# Pulvinar-cortical coupling in human epilepsy: spatiotemporal seizure dynamics with implications for neuromodulation

**DOI:** 10.64898/2026.11.27.769822

**Authors:** Yuduo Cai, Chaitanya Ganne, Stephen Harward, Gerald Grant, Derek Southwell, Shruti Agashe

**Affiliations:** Department of Biomedical Engineering, Duke University, NC, USA; Texas Institute of Restorative Neurotechnologies (TIRN), Department of Neurology, The University of Texas Health Science Center at Houston (UTHealth), Houston, TX; Department of Neurosurgery, Duke University, NC, USA; Department of Neurology, Duke University, NC, USA

**Keywords:** Thalamus, responsive neurostimulation, deep brain stimulation, stereo-EEG, intracranial EEG, drug resistant epilepsy

## Abstract

Despite growing interest in the pulvinar as a neuromodulation target, its dynamic engagement in human seizure networks remains poorly characterized. We analyzed simultaneous pulvinar and cortical stereo-electroencephalography from 15 participants and 30 seizures using time-resolved phase-amplitude coupling (tPAC). Ictal coupling between pulvinar theta-, alpha-, and beta-phase activity and cortical high-frequency activity increased significantly, with greater seizure-related increases in seizure-onset-zone (SOZ) cortex for alpha and beta. Pulvinar beta-phase coupling in SOZ cortex showed the earliest cohort-level departure from calibrated preictal fluctuations. SOZ coupling exhibited greater post-onset rise rates across all bands and earlier peaks for pulvinar theta- and alpha-phase coupling. Early-ictal pulvinar theta- and alpha-phase coupling was numerically strongest in parahippocampal regions, and beta-phase coupling in inferior occipital regions. Temporal cortex exhibited the earliest regional departure in pulvinar beta-phase coupling, whereas occipital cortex reached peak coupling fastest across all bands. These findings suggest frequency-dependent and anatomically differentiated pulvinar engagement during human seizures.

## 1. Background

Closed-loop and open-loop intracranial neuromodulation have emerged as important therapies for drug-resistant epilepsy, with thalamic targets increasingly recognized as key modulatory nodes within distributed seizure networks^1–3^. Among candidate targets, the pulvinar nucleus has drawn increasing interest because of imaging evidence for differentiated anatomical and functional connections with occipital, temporal, and parietal cortical areas^4,5^ as well as broader functional coactivation patterns identified by meta-analysis^6^. Emerging clinical studies with small cohorts have also reported potential benefits of pulvinar stimulation in selected patients with drug-resistant epilepsy^7–12^. Yet the extent to which the pulvinar is engaged during various phases of seizures, from onset to evolution remains incompletely defined, largely because direct peri-ictal recordings from the pulvinar are rarely available^13–16^.

Stereo-electroencephalography (sEEG) provides a unique opportunity to sample the pulvinar concurrently with cortical seizure-onset zones (SOZs), enabling direct interrogation of thalamocortical interactions during human seizures^17^. Whereas cortical seizures often begin with fast oscillatory activity, thalamic ictal activity may exhibit distinct spectral characteristics and morphologies, with thalamic recruitment occurring synchronously with cortical seizure onset or after a variable delay^17–19^. Characterizing the spatial and temporal dynamics of pulvinar-cortical coupling, including its relationship to seizure-onset regions, timing of engagement, and extent across the ictal network, may identify physiological signatures of pulvinar engagement that could ultimately inform thalamic targeting, detection and closed-loop neuromodulation therapies in epilepsy.

Phase-amplitude coupling (PAC) is a form of cross-frequency coupling that quantifies the statistical dependence between the phase of a low-frequency oscillation and the amplitude of a higher-frequency component of neural activity^20,21^. Physiologically, PAC may reflect cross-timescale organization of neuronal activity, whereby slower oscillations modulate fluctuations in neuronal excitability and thereby influence the expression and timing of faster population activity^22,23^. Intracranial EEG studies have associated PAC measures with the distribution of pathological tissue and postoperative outcome in epilepsy^24^, localization of epileptogenic zone or seizure-onset regions^25,26^, and temporal changes during seizures^27^. The reliability of PAC estimates depends on the estimation method and the amount of data available, with shorter analysis windows improving temporal resolution at the expense of estimation stability^21,28^. Although sliding-window approaches have been used to characterize peri-ictal PAC dynamics in human intracranial EEG^29,30^, resolving rapid changes in coupling during seizure initiation and propagation remains challenging^31,32^. The time-resolved phase-amplitude coupling (tPAC) estimator developed by Samiee and Baillet (2017) offers an approach to characterizing rapid coupling dynamics with its performance evaluated in simulations and rat entorhinal cortex recordings^28^. This approach enables finer temporal characterization of coupling while retaining frequency-specific information.

Applying tPAC to simultaneous pulvinar and cortical stereo-EEG recordings enables characterization of coupling between the phase of multiple frequency bands in the pulvinar and the amplitude of high-frequency cortical activity, particularly within SOZs. Leveraging rare human pulvinar recordings, we investigate the spatial and temporal evolution of pulvinar-cortical coupling across different phases of seizures and seizure networks, with the goal of understanding pulvinar engagement during ictal activity and its potential implications as a neuromodulation target in epilepsy.

## 2. Methods

### 2.1 Patient selection

Fifteen patients with drug-resistant epilepsy who underwent sEEG as part of routine presurgical evaluation at Duke University were retrospectively identified for inclusion in the study. All patients were implanted according to clinical indications, with electrode trajectories and target selection determined independently by the treating multidisciplinary epilepsy team. Inclusion criteria were: (1) capture of at least one electrographic seizure during the sEEG monitoring admission, and (2) sampling of the pulvinar nucleus by one or more implanted contacts. Intracranial recordings were obtained using clinically implanted stereotactic depth electrodes (DIXI Medical or PMT Corporation) composed of cylindrical contacts (0.8 mm diameter, 2 mm contact length, 1.5 mm intercontact spacing). Across the cohort, 30 seizures were selected for analysis after confirmation of adequate signal quality, absence of artifact limiting interpretation, stable interictal baseline segments between seizures, and interpretable thalamic recordings.

### 2.2 Contact localization and preprocessing of sEEG

All imaging and electrode-localization procedures were performed using the Brainstorm sEEG processing pipeline^33^. The pre-implantation T1-weighted MRI served as the anatomical reference, and Montreal Neurological Institute (MNI) normalization was computed using SPM12^34^. Post-implantation CT images were co-registered and resliced to the T1-weighted MRI using SPM12, with registration visually verified on the overlaid images. A thresholded isosurface mesh was generated from the co-registered CT to distinguish the high-density electrode contacts from surrounding tissue. Electrodes were reconstructed by selecting the appropriate electrode model and contact count and defining the electrode tip and entry point on the co-registered images. Reconstructed contacts were visually inspected on the MRI slices and CT isosurface.

Cortical anatomical labels were assigned using the Automated Anatomical Labeling Atlas 3 (AAL3) MNI parcellation in Brainstorm^35^. Thalamic contacts were localized using the Krauth-Morel stereotactic atlas of the human thalamus^36,37^. For pulvinar subdivision analyses, atlas probabilities for the anterior (PuA), medial (PuM), lateral (PuL), and inferior (PuI) pulvinar were summed across the two contacts comprising each seed pair; the subdivision with the largest positive summed probability was assigned, whereas ties or the absence of positive probabilities were classified as unclassified. Atlas-based anatomical assignments were additionally visually reviewed for anatomical plausibility (Fig. 1).

**Figure 1.**
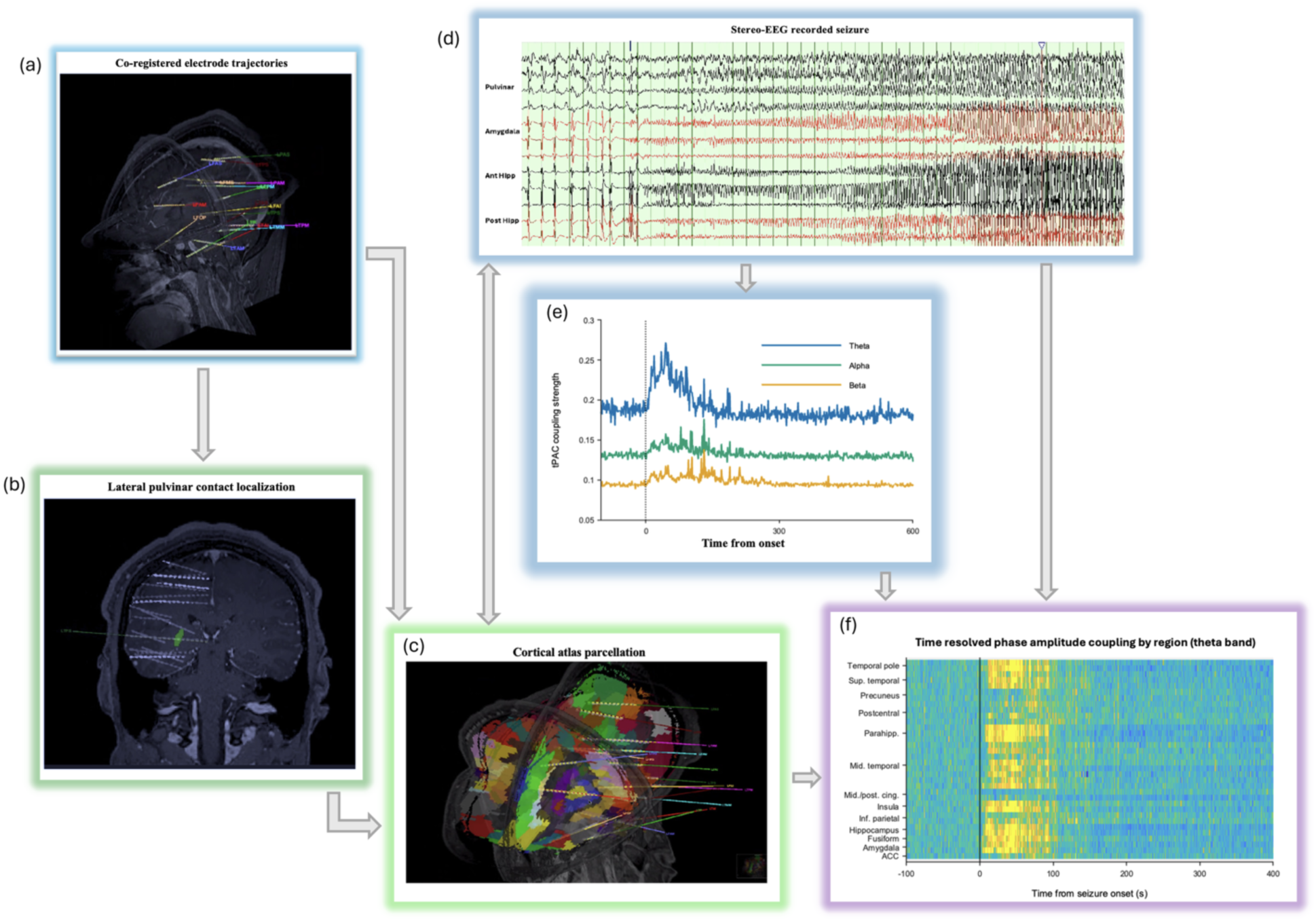
Electrode localization and time-resolved analysis of pulvinar-cortical coupling. **a**, Three-dimensional reconstruction of intracranial electrode trajectories co-registered to the participant’s anatomical MRI. **b,** Localization of pulvinar electrode contacts using the registered thalamic atlas. **c,** Electrode trajectories and contact locations overlaid on cortical anatomical parcellations used for regional assignment. **d,** Representative stereo-electroencephalography (sEEG) recording illustrating ictal activity in pulvinar and cortical seizure-onset channels. **e,** Corresponding time-resolved phase-amplitude coupling (tPAC) trajectories between pulvinar theta-, alpha-, and beta-band phase activity and cortical high-frequency amplitude (30-150 Hz). The vertical dashed line indicates electrographic seizure onset. **f,** Representative cortical channel-by-time heat map of baseline-relative change in pulvinar theta-phase coupling (ΔtPAC), with cortical targets grouped by anatomical region. Time is referenced to electrographic seizure onset, and ΔtPAC is expressed as percentage change from the preictal baseline.

**Figure 2.**
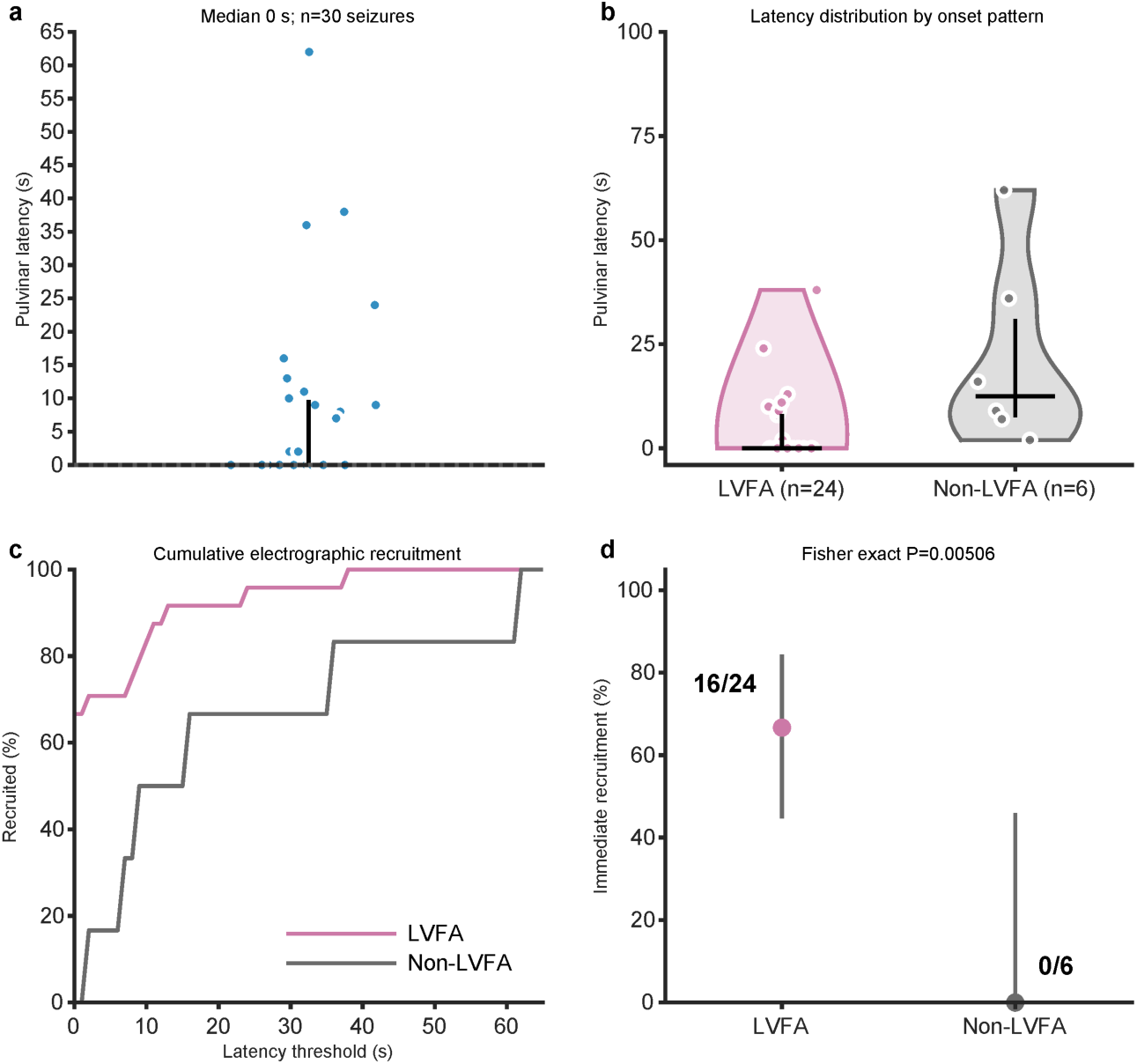
Electrographic recruitment of the pulvinar during seizures. **a**, Pulvinar recruitment latency relative to cortical electrographic seizure onset across 30 seizures from 15 participants (median, 0 s; interquartile range [IQR], 0-9.75 s; range, 0-62 s). **b,** Recruitment latency stratified by seizures with cortical low-voltage fast activity (LVFA; n = 24) and non-LVFA onset patterns (n = 6). Violin outlines represent kernel density estimates truncated to the observed range; black bars indicate median and IQR. **c,** Cumulative proportion of seizures showing electrographic pulvinar recruitment as a function of latency, stratified by LVFA status. Across all seizures, recruitment occurred at cortical onset in 16/30 (53.3%), within 5 s in 18/30 (60.0%), and within 10 s in 23/30 (76.7%). **d,** Proportion of seizures with immediate pulvinar recruitment in LVFA versus non-LVFA onset patterns (16/24 versus 0/6; two-sided Fisher’s exact P = 0.00506). Whiskers indicate unadjusted exact 95% binomial confidence intervals. Individual points in **a**, **b** represent seizures. All statistical comparisons were descriptive and unadjusted because individual participants contributed multiple seizures.

sEEG recordings were acquired using a Natus Medical Quantum LTM amplifier (Natus Medical, Pleasanton, CA, USA) at a sampling rate of 2048 Hz. Signals were band-pass filtered from 0.1-1000 Hz and notch filtered at 60 Hz. Recordings were then re-referenced to a bipolar montage to reduce volume conduction and far-field contamination, particularly from gray-matter sources detected by adjacent white-matter contacts^38,39^. The resulting recordings were visually inspected for artifacts and signal quality. Cortical contacts were classified as SOZ or non-SOZ based on clinical documentation, with these designations verified by a board-certified epileptologist (S.A.). For visual analysis, pulvinar recruitment timing was assessed from sEEG recordings using available clinical annotations and independently verified by a board-certified epileptologist (S.A.) relative to electrographic seizure onset.

For descriptive spatial visualization, each eligible cortical bipolar target was placed at the midpoint of its two contacts’ MNI coordinates and projected onto the nearest vertex, within 20 mm of the SPM12 canonical MNI cortical mesh (8,196 vertices); pulvinar seed contacts were not mapped. Mean ictal tPAC and ΔtPAC (Section 2.4) were rendered on the translucent mesh in left, right, anterior and posterior views with custom MATLAB scripts (R2021b), separately for theta-, alpha- and beta-phase coupling and for SOZ (Fig. 6) and non-SOZ (Supplementary Fig. 1) targets. Within each band and measure, SOZ and non-SOZ maps shared color limits were set to the 5th-95th percentiles of all mapped targets. These spatial maps were intended to illustrate the distribution of sampled coupling measures and were not used for anatomical interpolation or statistical inference.

### 2.3 Dynamic phase-amplitude coupling analysis

For each seizure, pulvinar bipolar pairs were used as low-frequency phase seeds and cortical bipolar pairs as high-frequency amplitude sources (Fig. 1d, e). Inter-regional tPAC was computed using the Samiee-Baillet estimator implemented in Brainstorm^28^. Pulvinar phase was evaluated separately in the theta (4-8 Hz), alpha (8-12 Hz), and beta (13-30 Hz) bands, while cortical high-frequency amplitude was evaluated over 30-150 Hz using a 1.1-s sliding window with 50% overlap and a 2-s filter margin. The algorithm identified a phase-frequency component within each specified pulvinar band and evaluated its coupling with cortical high-frequency amplitude at 20 linearly spaced center frequencies. Brainstorm linearly interpolated the output to a 0.275-s time grid. This interpolation increased the temporal sampling density without improving the independent temporal resolution of coupling estimation. Within each window, coupling strength, hereafter tPAC, was quantified as the length of the mean composite vector formed by the cortical high-frequency amplitude envelope and the pulvinar phase^22^, normalized by the root-mean-square amplitude as implemented in the Brainstorm tPAC estimator^28,40^:

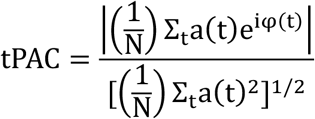

where φ(t) is the pulvinar phase, a(t) the cortical amplitude envelope and N the number of samples retained for coupling estimation, generally spanning complete phase cycles within the analysis window. tPAC is dimensionless and bounded between 0 and 1. For each specified pulvinar phase band and time window, coupling strength was summarized using Brainstorm’s maxPAC output, representing the maximum normalized coupling estimate across the evaluated cortical high-frequency amplitude range rather than the mean across that range. Of 267 declared seed-band maps, 225 maps from 75 seizure-specific seed pairs met seed-eligibility criteria. Seeds localized outside the pulvinar, within non-pulvinar structures, or within white matter were excluded; a stricter sensitivity analysis retained only seeds with good or acceptable localization. This mean-vector-length measure differs from the entropy-based modulation index described by Tort et al. (2010)^21^. Because absolute coupling values are estimator-specific, coupling strength is reported throughout as dimensionless tPAC.

### 2.4 Quantification of pulvinar-cortical tPAC measures

Coupling magnitude was quantified as mean tPAC over the specified analysis interval. Full-ictal analyses encompassed the interval from electrographic seizure onset to offset, whereas early-ictal analyses were restricted to the first third of each seizure, including regional coupling magnitude, peak latency, and maximum-velocity measures. Baseline-relative change (ΔtPAC) was calculated as the percentage change relative to the preictal baseline, using the absolute baseline value as the denominator (Fig. 1f). Absolute ictal-minus-baseline differences were reported as dimensionless tPAC differences, whereas differences in ΔtPAC were expressed in percentage points.

Peak tPAC was defined as the earliest maximum of a nominal 10-s moving-average trajectory within the specified ictal analysis interval. Time to peak tPAC was calculated from electrographic seizure onset to this maximum, using the full ictal interval for SOZ-non-SOZ comparisons and the early ictal third for regional analyses. Peak latencies were calculated for each eligible pulvinar seed-cortical target trajectory and averaged across seeds within each target. For paired SOZ-non-SOZ comparisons, target-level peak latencies were summarized as medians within seizures and subsequently across seizures within participants. Maximum velocity was defined separately as the largest numerical derivative of the smoothed tPAC trajectory, using nominal 10-s smoothing, with 5-s smoothing examined as a sensitivity analysis. Regional maximum-velocity analyses were restricted to the early ictal third.

Global and SOZ rise rates were calculated from ΔtPAC trajectories on a common 2-s grid following nominal 10-s smoothing. Rise rate was defined as the secant slope from the smoothed value at seizure onset to the maximum within the available 0-300-s post-onset window, expressed in percentage points s⁻¹. One seizure had less than 300 s of available post-onset recording. Because this window could extend beyond seizure offset, these rates were interpreted as post-onset rather than exclusively ictal changes. Global averages used available-channel denominators, excluding zones with entirely missing data from both the weighted sum and denominator. Smoothed time centers outside recorded support were excluded, and a maximum at onset yielded an undefined rise rate. Pulvinar subdivision-specific baseline-to-peak secant rates were calculated separately using the difference between mean preictal tPAC and peak ictal tPAC, divided by onset-to-peak latency, and expressed in tPAC s⁻¹, as detailed in Section 2.7.

### 2.5 Temporal evolution of pulvinar-cortical tPAC

To characterize the evolution of pulvinar-cortical tPAC across seizures, coupling trajectories were aligned to electrographic seizure onset, with baseline coupling calculated over the available preictal interval (−100 to −10 s; −54 to −10 s for one participant). The ictal period was divided into seizure-specific early, middle, and late thirds. Onset-aligned trajectories were resampled to a common 2-s grid spanning −100 to +600 s, with unavailable samples retained as missing. To account for differences in seizure duration, ictal trajectories were also normalized to 101 equally spaced positions spanning 0-100% of electrographic seizure duration^41^. When the first or last ictal sample center did not coincide exactly with seizure onset or offset, the nearest available ictal value was assigned to the corresponding 0% or 100% boundary to anchor the normalized trajectory. For the longer onset-aligned trajectories shown in Fig. 3a, participant-level trajectories were smoothed using a centered five-sample moving mean on the 2-s grid. Group-level trajectories were calculated across available participants, with each participant contributing equally, and are presented as mean ± SEM.

**Figure 3.**
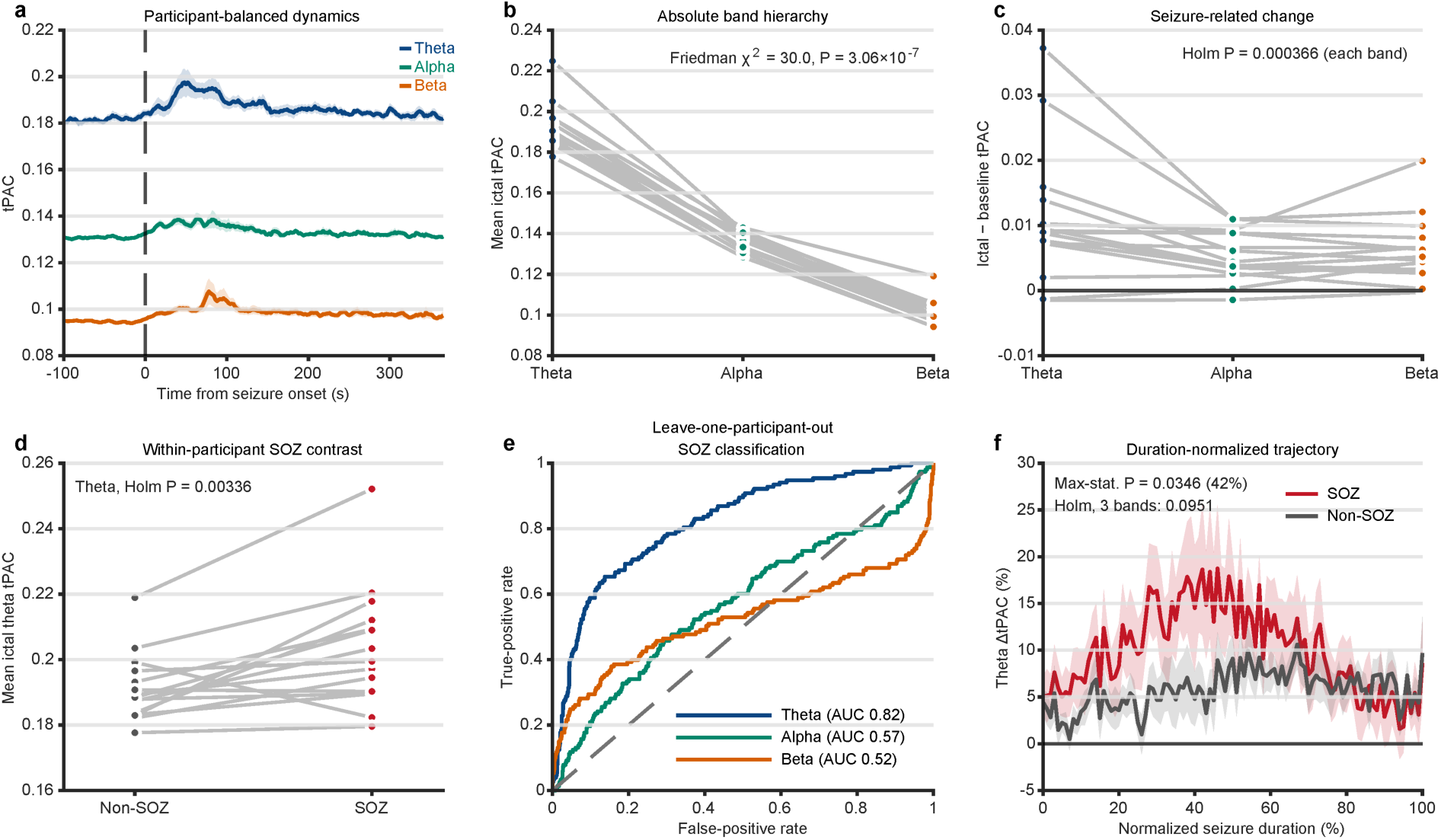
Ictal pulvinar-cortical coupling magnitude and SOZ-specific differences. **a**, Participant-balanced, seizure-onset-aligned trajectories of pulvinar theta-, alpha-, and beta-phase tPAC; shading indicates SEM across participants. **b,** Participant-level mean tPAC across phase bands over the full ictal interval. **c,** Participant-level absolute ictal-minus-baseline tPAC differences across phase bands. **d,** Paired comparison of full-ictal theta-phase tPAC between seizure-onset-zone (SOZ) and non-SOZ contacts. Grey lines in **b-d** connect observations from the same participant (n = 15). **e,** Pooled receiver-operating-characteristic (ROC) curves generated from out-of-fold predictions using leave-one-participant-out logistic regression with baseline and ictal-minus-baseline tPAC features. Each contact contributed to the pooled ROC analysis; the dashed diagonal indicates chance-level discrimination. **f,** Duration-normalized pulvinar theta-phase ΔtPAC in SOZ and non-SOZ contacts across 0-100% of electrographic seizure duration; shading indicates participant-level uncertainty. The maximum SOZ-non-SOZ separation occurred at 42% of seizure duration (13.14 percentage points; temporal family-wise P = 0.0346) but did not survive additional Holm correction across the three prespecified phase bands (adjusted P = 0.0951). Statistical comparisons in **b-d** used participant-level Friedman or paired Wilcoxon signed-rank tests, with Holm correction as appropriate. ΔtPAC denotes percentage change relative to preictal baseline; absolute tPAC differences are dimensionless.

### 2.6 Focused peri-onset analysis of pulvinar-cortical tPAC

Focused peri-onset analyses evaluated the earliest detectable departure of pulvinar-cortical tPAC from baseline. These analyses included 14 participants (28 seizures), excluding recordings with insufficient baseline data following nested calibration or incomplete 10-s ictal coverage. Native tPAC estimates were recovered from interpolated maps at nominal 0.55-s intervals. At each shared analysis time point, the most recent eligible estimate was used, allowing a maximum carry-forward of 0.55 s. Windows overlapping seizure onset were excluded, yielding an earliest evaluable post-onset time of 1.65 s. A phase-support sensitivity analysis required each native window center and its surrounding 2.55-s interval (2-s phase-support margin plus 0.55-s half-window) to remain entirely within the preictal or ictal period, shifting the earliest evaluable post-onset time to 3.85 s. These limits reflect retrospective analytical constraints, do not eliminate dependencies introduced by whole-record filtering and Hilbert transforms, and should not be interpreted as real-time detection latencies.

For peri-onset analyses, pulvinar seeds were averaged within each cortical target and seizure without selection based on ictal coupling. Each target-level trajectory was standardized to its preictal mean and sample standard deviation over native window centers from −100 to −13.2 s relative to seizure onset. Targets were averaged within seizures, seizures within participants, and participants equally. SOZ-non-SOZ and regional contrasts were calculated within seizures before participant-level aggregation. Recent-preictal mean levels and slopes were jointly tested across nine SOZ, non-SOZ, and direct SOZ-non-SOZ trajectories (18 components). Reference drift was evaluated separately using late-minus-early reference-third differences and slopes across the full reference interval in a separate 18-component participant family. To account for recent preictal trends, linear forecasts fitted to each seizure’s pre-onset values at their actual native window centers were subtracted from post-onset values. Constant recent-preictal mean forecasts and the phase-support restriction were evaluated as separate sensitivity analyses. A secondary contact-level multiscale detector used 1, 2, 4, 8, and 16 native estimates and an independent −100 to −10-s reference interval. Detection fraction, non-detection-inclusive restricted time, and latency conditional on detections in both SOZ and non-SOZ contacts were treated as distinct endpoints.

### 2.7 Cortical regional and pulvinar subdivision analyses

Regional analyses were performed at two anatomical resolutions comprising seven broad anatomical groups (six cortical and one thalamic) and 25 finer subgroups (Supplementary Table 1). Both contacts of a bipolar target were required to have valid atlas labels and belong to the same anatomical group or subgroup, as appropriate. Additional parcel-level analyses required both contacts to share the same AAL3 base label. Temporal subgroups included mesial temporal, lateral temporal neocortical, ventral temporal/occipitotemporal, and auditory cortical regions, whereas occipital subgroups included primary visual and visual association regions. The complete anatomical grouping scheme is provided in Supplementary Table 1. Cerebellar, basal ganglia, white-matter, and unassigned targets were excluded, while thalamic targets were retained as a separate broad group. Hemispheres were pooled. Targets sharing a physical contact with the current pulvinar seed were excluded, and cortical targets containing any eligible pulvinar seed contact within that seizure were additionally excluded. Eighteen duplicated seed-channel-band time series were removed before aggregation. Broad-region estimates were calculated directly from eligible targets rather than subgroup-level averages.

For regional analyses, eligible pulvinar seed estimates were averaged within each cortical target. Coupling magnitude and time to peak tPAC were calculated over the specified intervals (Section 2.4), with target-level estimates aggregated within seizures and subsequently within participants. Means were reported for coupling magnitude and cohort medians for temporal endpoints. Regional peri-onset departures were evaluated separately using the focused baseline-departure analysis (Section 2.6), with participant-balanced trajectories and region-versus-rest contrasts restricted to cortical anatomical groups.

For pulvinar subdivision analyses, seed-target coupling was summarized within seizure, cortical region, frequency band, and subdivision before participant-level aggregation. Multiple eligible seed pairs within the same subdivision were averaged before aggregation. Subdivision baseline-to-ictal-peak secant rates were calculated as the difference between the smoothed ictal maximum and mean preictal baseline, divided by onset-to-peak latency, expressed in tPAC s⁻¹. Rates were calculated only when the selected maximum had full nominal 10-s smoothing support; otherwise, they were treated as missing without selecting an alternative peak. These rates differed from the baseline-relative global/SOZ rise rates (Section 2.4). PuM-PuL comparisons were descriptive because fewer than six matched participants were available.

Focused peri-onset analyses were not stratified by pulvinar subdivision because eligible seeds were averaged within targets before trajectory analysis. Magnitude and rate analyses required at least three eligible participants; conditional latency estimates with fewer participants were flagged separately. All analyses remained frequency-band specific.

### 2.8 Statistical analysis

The participant was the primary unit of statistical inference unless otherwise specified.

All tests were two-sided, with P < 0.05 considered statistically significant after multiplicity correction where applicable. Within-participant comparisons used Wilcoxon signed-rank tests, whereas comparisons across three frequency bands or ictal epochs used Friedman tests. For Wilcoxon analyses, W denotes the smaller signed-rank sum. Primary paired effects were expressed as median within-participant differences, with signed rank-biserial correlations as effect sizes^42^. For the primary paired effects in Fig. 3, 95% percentile confidence intervals were obtained using 5,000 participant-level bootstrap resamples^43^. Secondary contact-level effect estimates and intervals used the endpoint-specific definitions below. Holm correction was applied within prespecified frequency-band and contrast families^44^. Nonsignificant findings were not interpreted as evidence of equivalence.

Focused peri-onset analyses used synchronized baseline pseudo-epochs to construct two-sided maximum-statistic bounds, corrected across trajectories and time points. Distant-reference SOZ analyses included separate families of six absolute trajectories and nine paired zone/frequency-band contrasts, each corrected across 34 pre-and post-onset endpoints. Regional and region-versus-rest analyses used separate 18-curve families, while local-linear residual analyses used a nine-curve family across 16 post-onset endpoints. Crossing a corrected baseline envelope indicated departure from preictal fluctuations rather than biological coupling onset. Participant-level contrasts and local-linear residuals were tested separately using exact synchronized whole-participant sign permutations, with family-wise correction across trajectories and time^45,46^. Descriptive pointwise 95% percentile intervals were obtained using 5,000 participant-level bootstrap resamples for distant-reference trajectories and 2,000 for local-linear residuals, separately from the corrected baseline-null bounds.

In the secondary contact-level peri-onset analysis, effects were the mean of within-participant paired differences, with pointwise 95% percentile intervals from 10,000 participant-level bootstrap resamples. Paired comparisons used exact two-sided signed-rank inference. For SOZ comparisons, Holm correction covered six primary tests (three bands × detection fraction and restricted time at 10 s), six early-detection tests (three bands × detection fraction at 2 and 5 s), and a separate three-band conditional-latency family. Each horizon included seizures with complete follow-up through that endpoint. Conditional latencies were not interpreted as unbiased SOZ-non-SOZ onset differences.

Duration-normalized SOZ-non-SOZ trajectories were evaluated using 10,000 participant-level sign permutations, with the maximum absolute mean contrast across 101 normalized seizure-time positions controlling temporal family-wise error. Early, middle, and late ictal summaries corresponded to 0-33%, 34-66%, and 67-100% of seizure duration, respectively, with Holm correction across prespecified band-by-epoch comparisons. Global rise rates were compared across frequency bands using Friedman tests, while paired SOZ-non-SOZ rise rates were compared using Wilcoxon signed-rank tests with Holm correction across bands. Exploratory regional comparisons used paired two-sided nonparametric tests, requiring at least six matched participants, with Holm correction within prespecified comparison families. Regional ictal-phase analyses compared ictal thirds within each frequency band, with correction applied within band. Pulvinar subdivision comparisons were descriptive because fewer than six matched participants were available. LVFA and seizure-level clinical timing comparisons were also considered descriptive because multiple seizures were contributed by individual participants.

The internal SOZ classifier was evaluated using leave-one-participant-out logistic regression with baseline tPAC and ictal-minus-baseline tPAC as features. Feature scaling and model fitting were performed using training participants only. Classification performance was summarized using both pooled out-of-fold AUC and the median of within-participant AUCs to distinguish overall held-out discrimination from contact-level ranking within individual participants. Because SOZ labels and ictal features were derived from the same recordings, classification performance reflected internal separability rather than independent diagnostic accuracy. Statistical analyses were performed in Python using NumPy^47^, SciPy^48^, pandas^49^, statsmodels^50^, and scikit-learn^51^; tPAC maps were generated in MATLAB using Brainstorm^33^.

## 3. Results

### 3.1 Visual sEEG analysis demonstrates early pulvinar recruitment

The final analytic cohort comprised 15 participants and 30 seizures. Median seizure duration was 94.5 s (interquartile range, 54.75-148.0; range, 7-1010 s). Twenty-four seizures had cortical low-voltage fast activity (LVFA^52^) while six had non-LVFA onset patterns (Fig. 2). Temporal regions were involved at onset in 25 of 30 seizures (83.3%), and occipital regions in 10 of 30 seizures (33.3%). Across subgroups, mesial temporal onset was most frequent (15/30, 50.0%), followed by lateral temporal onset (12/30, 40.0%). Ventral temporal/occipitotemporal involvement was present in 6 seizures (20.0%), inferior occipital gyrus involvement in 6 (20.0%), and lingual gyrus involvement in 4 (13.3%). Insular-opercular onset was present in 3 seizures (10.0%). No seizures demonstrated onset within the predefined parietal regions. Because individual seizures could involve more than one anatomical region or subgroup at onset, these categories were not mutually exclusive. Onset was confined to temporal regions in 18 seizures (60.0%) and to occipital regions in 5 (16.7%); the remaining 7 seizures (23.3%) involved more than one broad region (temporal and occipital in 5, one of which also involved the insula, and temporal and insular in 2). Among the 25 seizures with temporal involvement, 6 involved more than one temporal subgroup.

Across 30 seizures, the median visually determined pulvinar recruitment latency relative to cortical seizure onset was 0 s (IQR, 0-9.75 s; range, 0-62 s; Fig. 2a; stratified by onset pattern in Fig. 2b). Using cumulative latency thresholds, pulvinar recruitment coincided with cortical onset in 16 of 30 seizures (53.3%), occurred within 5 s in 18 of 30 (60.0%), and within 10 s in 23 of 30 (76.7%; Fig. 2c). Immediate recruitment was visually observed in 16 of 24 seizures with low-voltage fast activity (LVFA) onset and in none of the 6 non-LVFA seizures (two-sided Fisher exact P = 0.00506; Fig. 2d). These seizure-level comparisons were considered descriptive because multiple seizures were contributed by individual participants.

### 3.2 Pulvinar-cortical coupling increases during seizures and is greater in SOZ cortex

Across the full ictal interval, pulvinar-cortical tPAC increased significantly relative to the preictal baseline across pulvinar theta-, alpha-, and beta-phase frequencies, with median participant-level ictal-minus-baseline tPAC differences of 0.0089, 0.0044, and 0.0052 (median relative increases, 5.0%, 3.3%, and 5.5%), respectively (W = 3, 2, and 1; all Holm-adjusted P = 0.000366; rank-biserial correlations = 0.950, 0.967, and 0.983; n = 15; Fig. 3c). Participant-balanced, seizure-onset-aligned trajectories demonstrated distinct frequency-dependent temporal profiles, with a sustained post-onset increase in theta-phase coupling, a smaller increase in alpha-phase coupling, and a sharper beta-phase increase followed by decline (Fig. 3a). Theta-phase coupling showed the highest absolute magnitude across the displayed onset-aligned interval, consistent with the participant-level mean ictal hierarchy (Fig. 3a, b). Mean tPAC over the full ictal interval differed across phase-frequency bands (Friedman χ²(2) = 30.0, P = 3.06 × 10⁻⁷; Kendall’s W = 1.00), with theta exceeding alpha by a median paired difference of 0.0537, theta exceeding beta by 0.0900, and alpha exceeding beta by 0.0354 (all paired W = 0; Holm-adjusted P = 0.000183; Fig. 3b).

Mean ictal pulvinar-cortical tPAC was greater in SOZ than non-SOZ contacts across all three phase-frequency bands. Participant median tPAC values for SOZ versus non-SOZ contacts were 0.19945 versus 0.18843 for theta, 0.14162 versus 0.13628 for alpha, and 0.10792 versus 0.09831 for beta. Median paired SOZ-non-SOZ differences were 0.01230 for theta (95% CI, 0.00193-0.02057; W = 11; Holm-adjusted P = 0.00336), 0.00478 for alpha (95% CI, 0.00268-0.00992; W = 6; adjusted P = 0.00171), and 0.00631 for beta (95% CI, 0.00260-0.01914; W = 3; adjusted P = 0.000916; n = 15; Fig. 3d shows theta). Because SOZ-non-SOZ differences were also present at baseline, absolute ictal contrasts were complemented by difference-in-differences analyses. The seizure-related increase in coupling was greater in SOZ than non-SOZ contacts for alpha (median paired difference, 0.00484; W = 16; adjusted P = 0.0205) and beta (0.00575; W = 8; adjusted P = 0.00458), but not theta (0.00792; W = 27; adjusted P = 0.0637).

Exploratory leave-one-participant-out classification using baseline and ictal-minus-baseline tPAC features demonstrated the strongest pooled discrimination for theta-phase coupling (AUC = 0.821), whereas alpha- and beta-phase coupling showed near-chance pooled performance (AUC = 0.573 and 0.523, respectively; 15 participants, 1,339 contacts, 153 of them SOZ; Fig. 3e). Median within-participant AUCs were higher across all three bands (0.888, 0.841, and 0.828, respectively), suggesting that within-participant SOZ-non-SOZ contact ranking may be more consistent than discrimination across pooled held-out contacts. These analyses reflect internal classification performance and do not establish independent diagnostic accuracy.

To examine coupling differences across seizure evolution, each ictal interval was normalized to 0-100% of its electrographic duration. The largest theta-phase SOZ-non-SOZ separation occurred at 42% of seizure duration (13.14 percentage points; 10,000-sign-flip maximum-statistic family-wise P = 0.0346; Fig. 3f), although this time-local effect did not survive Holm correction across the three phase bands (adjusted P = 0.0951) or restriction to seeds with good or acceptable localization (P = 0.27; 12 participants). The time-averaged SOZ-non-SOZ difference was also not statistically significant (median across participants, 4.26 percentage points; 95% bootstrap interval, −0.55 to 9.23; P = 0.0637).

### 3.3 Temporal dynamics of pulvinar-cortical coupling differentiate SOZ from non-SOZ contacts

Focused peri-onset analysis of 28 seizures from 14 participants identified the earliest post-onset cohort-level departure from the preictal reference in pulvinar beta-phase coupling to SOZ cortex. Pulvinar beta-phase coupling in SOZ cortex crossed the corrected distant-reference envelope at the first evaluable post-onset endpoint (1.65 s), followed by alpha-phase coupling at 3.85 s, whereas theta-phase coupling showed no departure through 9.90 s. No pulvinar phase-frequency band showed a corresponding departure in non-SOZ cortex during this interval (Fig. 4). Pulvinar beta-phase coupling in SOZ cortex also crossed the corrected baseline envelope once before seizure onset (−7.70 s; baseline-family P = 0.029), consistent with the recent-preictal elevation described in the Supplementary Results.

**Figure 4.**
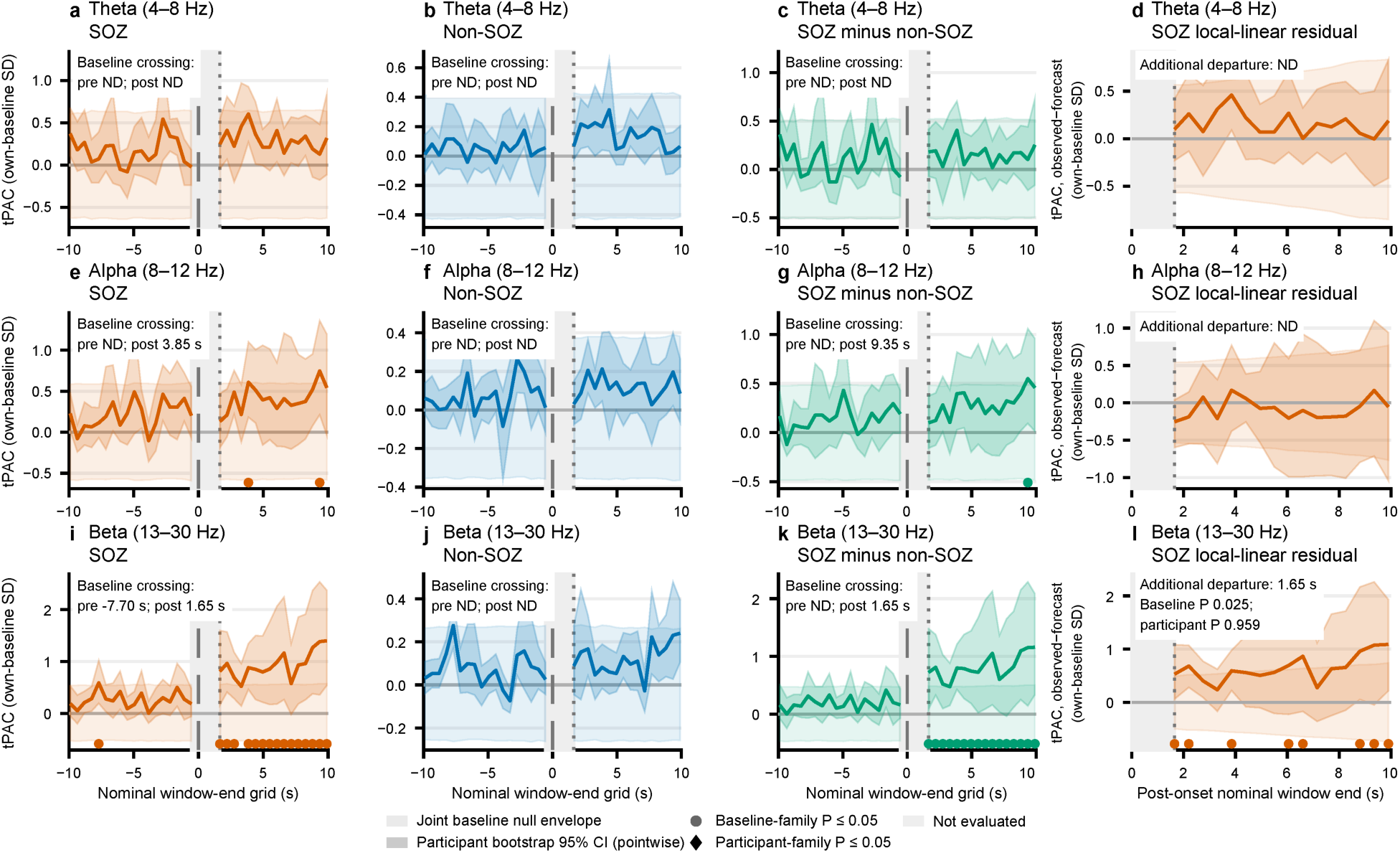
Early peri-onset pulvinar-cortical coupling dynamics in SOZ and non-SOZ cortex. Time-resolved pulvinar theta-, alpha-, and beta-phase coupling to cortical high-frequency activity was examined relative to electrographic seizure onset in 14 participants (28 seizures). a-c, e-g, i-k, Standardized coupling trajectories in seizure-onset-zone (SOZ) cortex, non-SOZ cortex, and their direct difference, respectively, for theta-(a-d), alpha-(e-h), and beta-phase (i-l) coupling. d, h, l, SOZ trajectories after subtraction of linear forecasts fitted to recent preictal activity. Pale envelopes indicate corrected baseline-null bounds, darker shading represents pointwise 95% participant-bootstrap intervals, and filled markers indicate departures beyond the corrected baseline envelope. Pulvinar beta-phase coupling in SOZ cortex showed the earliest post-onset departure at 1.65 s, followed by alpha-phase coupling at 3.85 s; theta-phase coupling and all non-SOZ bands showed no departure through 9.90 s. The baseline-referenced SOZ-non-SOZ beta-phase difference also departed at 1.65 s, with the early beta-phase departure persisting after recent-preictal detrending. Detection times represent analysis-dependent window endpoints rather than biological onset latencies.

In the same distant-reference analysis, the SOZ-non-SOZ contrast in pulvinar beta-phase coupling departed from its corrected preictal reference envelope at 1.65 s (baseline-family-adjusted P = 0.0006), whereas the corresponding alpha-phase contrast first departed at 9.35 s. These findings indicate changes in the baseline-referenced SOZ-non-SOZ coupling contrast rather than significant absolute post-onset differences between the two cortical groups. Indeed, no direct post-onset participant-level SOZ-non-SOZ contrast survived family-wise correction (minimum adjusted P = 0.251). The early beta-phase effect was heterogeneous: three participants accounted for 83.8% of the signed SOZ response and 89.6% of the SOZ-non-SOZ contrast at the first evaluable endpoint. Excluding these participants delayed the respective distant-reference departures to 4.40 and 3.85 s. The early pulvinar beta-phase coupling departure persisted after adjustment for recent-preictal trends and under the more conservative phase-support restriction (Supplementary Results).

Post-onset rise rates of pulvinar-cortical ΔtPAC further differentiated SOZ from non-SOZ contacts. Mean global rise rates, calculated over the available 0-300-s post-onset window, were 0.343, 0.269, and 0.361 percentage points s⁻¹ for pulvinar theta-, alpha-, and beta-phase coupling, respectively, with no significant difference across bands (Friedman χ²(2) = 1.60, P = 0.4493; n = 15; Fig. 5a). When compared by cortical zones, rise rates were significantly greater in SOZ than non-SOZ contacts across all three pulvinar phase bands. Mean SOZ versus non-SOZ rise rates were 0.715 versus 0.286 percentage points s⁻¹ for theta (median paired difference, 0.358; W = 7; Holm-adjusted P = 0.00122), 1.158 versus 0.221 for alpha (difference, 0.699; W = 3; adjusted P = 0.000916), and 1.300 versus 0.320 for beta (difference, 0.574; W = 3; adjusted P = 0.00122; Fig. 5b). Paired rise-rate analyses included 15 participants for theta and alpha and 14 for beta.

**Figure 5.**
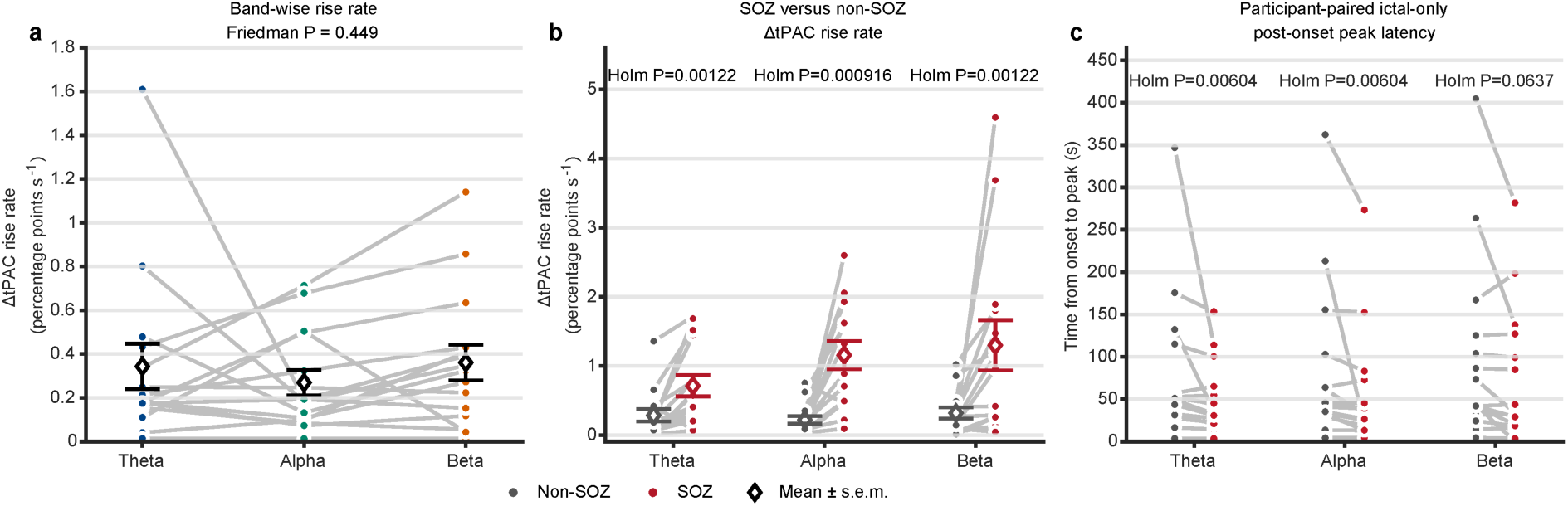
Rise rate and time to peak pulvinar-cortical tPAC distinguish SOZ from non-SOZ cortex. **a**, Participant-level global rise rates of pulvinar theta-, alpha-, and beta-phase ΔtPAC over the available 0-300-s post-onset window, including postictal activity. **b,** Paired comparison of post-onset ΔtPAC rise rates between seizure-onset-zone (SOZ) and non-SOZ cortical contacts across phase bands. **c,** Paired comparison of time to peak tPAC between SOZ and non-SOZ contacts, with peak searches restricted to the full electrographic ictal interval (onset to offset). Grey lines connect paired participant values. Rise rates were greater in SOZ than non-SOZ contacts across all three phase bands, whereas peak latency was shorter in SOZ for theta and alpha, but not beta, after correction. Global rise rates were compared using a Friedman test; paired SOZ-non-SOZ comparisons used two-sided Wilcoxon signed-rank tests with Holm correction across bands. Rise rates are expressed in percentage points s⁻¹. Paired rise-rate analyses included 15, 15, and 14 participants for theta, alpha, and beta, respectively; peak-latency analyses included 15 participants per band.

**Figure 6.**
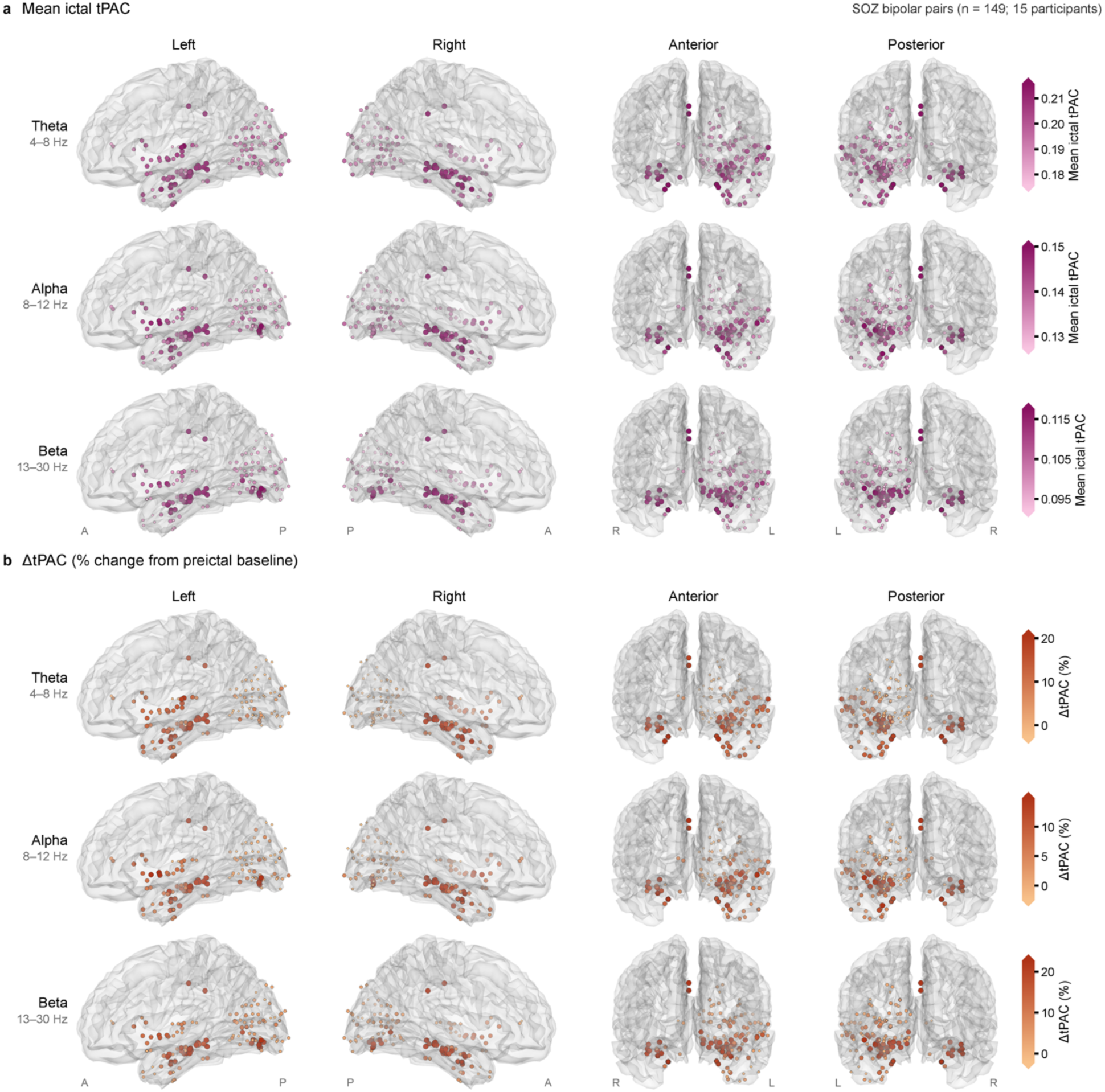
Spatial distribution of mean ictal tPAC and ΔtPAC across sampled SOZ cortex. Glass-brain maps showing pulvinar theta-(4-8 Hz), alpha-(8-12 Hz), and beta-phase (13-30 Hz) coupling across sampled seizure-onset-zone (SOZ) cortical bipolar targets, projected onto a standardized Montreal Neurological Institute (MNI) cortical surface in left, right, anterior, and posterior views. **a,** Mean tPAC over the full electrographic ictal interval. **b,** Baseline-relative change in tPAC (ΔtPAC), expressed as percentage change from the preictal baseline (−100 to −10 s). Each marker represents a cortical bipolar target projected from its midpoint coordinates onto the cortical surface, with marker color and size reflecting coupling magnitude or change. Color scales were matched between SOZ and non-SOZ maps (Supplementary Fig. 1) within each frequency band and measure. These maps are descriptive, reflect uneven clinical sampling, and were not used for anatomical interpolation or statistical inference.

Within the full electrographic ictal interval, pulvinar-cortical tPAC reached its peak earlier after seizure onset in SOZ than non-SOZ contacts for theta- and alpha-phase coupling. Participant median times to peak tPAC in SOZ versus non-SOZ contacts were 35.9 versus 51.2 s for theta, 32.9 versus 45.7 s for alpha, and 33.6 versus 43.1 s for beta. Median paired SOZ-non-SOZ differences in peak latency were −10.4 s for theta (W = 9; Holm-adjusted P = 0.00604), −8.04 s for alpha (W = 9; adjusted P = 0.00604), and −4.95 s for beta (W = 27; adjusted P = 0.0637; n = 15; Fig. 5c). In the stricter seed-localization subset of 12 participants and 23 seizures, only the alpha ictal-only peak-latency contrast remained significant after Holm correction (adjusted P = 0.0103).

### 3.4 Pulvinar-cortical coupling evolves dynamically across the ictal period

Participant-balanced absolute pulvinar-cortical coupling varied numerically across early, middle, and late ictal thirds, without significant differences across epochs in any pulvinar phase-frequency band. Mean tPAC values across early, middle, and late thirds were 0.18949, 0.19456, and 0.19225 for theta; 0.13509, 0.13732, and 0.13567 for alpha; and 0.09850, 0.09963, and 0.10440 for beta. Three-epoch Friedman tests were nonsignificant for theta (χ²(2) = 0.933; Holm-adjusted P = 0.6271), alpha (χ²(2) = 4.933; adjusted P = 0.2546), and beta (χ²(2) = 3.333; adjusted P = 0.3778; n = 15), indicating no significant differences in absolute coupling magnitude across ictal thirds.

Exploratory duration-normalized trajectories suggested temporal variation in SOZ-non-SOZ coupling differences (Fig. 3f). For pulvinar theta-phase coupling, the SOZ trajectory increased during the early portion of the seizure, reached its greatest elevation during the middle third, and declined toward ictal offset, whereas the non-SOZ trajectory showed a smaller numerical increase. Mean ΔtPAC values across early, middle, and late thirds were 9.35%, 14.36%, and 6.45% in SOZ contacts, compared with 3.99%, 6.94%, and 6.38% in non-SOZ contacts. The largest time-local theta-phase separation occurred at 42% of electrographic seizure duration (13.14 percentage points; Fig. 3f). Band-specific analyses identified time-local SOZ-non-SOZ coupling differences across all three prespecified pulvinar phase-frequency bands, with temporal maximum-statistic family-wise P values of 0.0346, 0.0432, and 0.0317 for theta, alpha, and beta, respectively. These findings did not survive additional Holm correction across the three bands (adjusted P = 0.0951 for each).

Third-specific analyses evaluated SOZ-non-SOZ differences within each ictal third. Theta differences were not significant within the early, middle, or late thirds after correction (adjusted P = 0.1293, 0.1055, and 1.000, respectively). In contrast, early-ictal SOZ-non-SOZ differences remained significant for alpha, with mean ΔtPAC values of 9.24% in SOZ versus 2.87% in non-SOZ contacts (median paired difference, 7.19 percentage points; W = 13; adjusted P = 0.04297), and for beta, with corresponding values of 15.05% versus 3.38% (median paired difference, 8.97 percentage points; W = 8; adjusted P = 0.01373). No other band-by-third comparison survived correction. In the stricter seed-localization subset (12 participants, 23 seizures), neither early-ictal contrast remained significant (adjusted P = 0.21 for alpha and 0.15 for beta).

### 3.5 Regional and cortical subgroup variation in pulvinar-cortical coupling

Descriptive contact-level glass-brain maps illustrated the distribution of full-ictal coupling measures across sampled SOZ sites (Fig. 6), with corresponding non-SOZ maps shown on matched color scales in Supplementary Fig. 1. Mean ictal tPAC was distributed across sampled cortical territories in all three phase bands, while ΔtPAC maps illustrated the spatial distribution of seizure-related coupling changes. These maps were descriptive and were not used to infer statistical differences between cortical regions.

During the early ictal third, temporal targets showed the numerically greatest broad-region theta- and alpha-phase coupling (mean tPAC, 0.1924 and 0.1372; n = 15), whereas occipital targets showed the greatest beta-phase coupling (0.1012; n = 8). Temporal-occipital differences were not significant in any band (Holm-adjusted P = 0.94). In matched comparisons, temporal alpha coupling exceeded parietal and frontal coupling (Holm-adjusted P = 0.0044 and 0.031, respectively). No other broad-region magnitude comparison established a consistent regional hierarchy after correction. Thalamic targets, analyzed as a separate group, showed early-ictal coupling within the range of the cortical groups (0.1889, 0.1353, and 0.0974 for theta, alpha, and beta; n = 15).

Cortical subgroup analyses further characterized these early-ictal patterns (Fig. 7a). Within the temporal group, mesial temporal targets showed numerically greater theta-, alpha-, and beta-phase coupling than lateral temporal neocortical targets (0.1996 versus 0.1895, 0.1415 versus 0.1357, and 0.1052 versus 0.0970, respectively; Fig. 7a). Primary visual and visual-association subgroups also showed descriptive differences in coupling magnitude, without significant subgroup differences after correction. Early-ictal absolute tPAC changes relative to baseline also varied across cortical subgroups (Fig. 7f). Mesial temporal cortex showed prominent theta-, alpha-, and beta-phase increases (0.018, 0.011, and 0.010 tPAC, respectively), with a similarly elevated theta-phase change in ventral temporal/occipitotemporal cortex (0.016 tPAC). In contrast, primary visual cortex showed smaller changes across all three bands (0.001, 0.001, and 0.003 tPAC), whereas visual association cortex showed a greater beta-phase increase (0.006 tPAC). At the finer AAL3 parcel level, parahippocampal cortex showed the greatest theta- and alpha-phase coupling (0.2022 and 0.1446 tPAC; n = 11), whereas inferior occipital cortex showed the greatest beta-phase coupling (0.1081 tPAC; n = 4) among cortical parcels represented in at least three participants. These descriptive findings highlight regional heterogeneity in both coupling magnitude and seizure-related changes.

**Figure 7.**
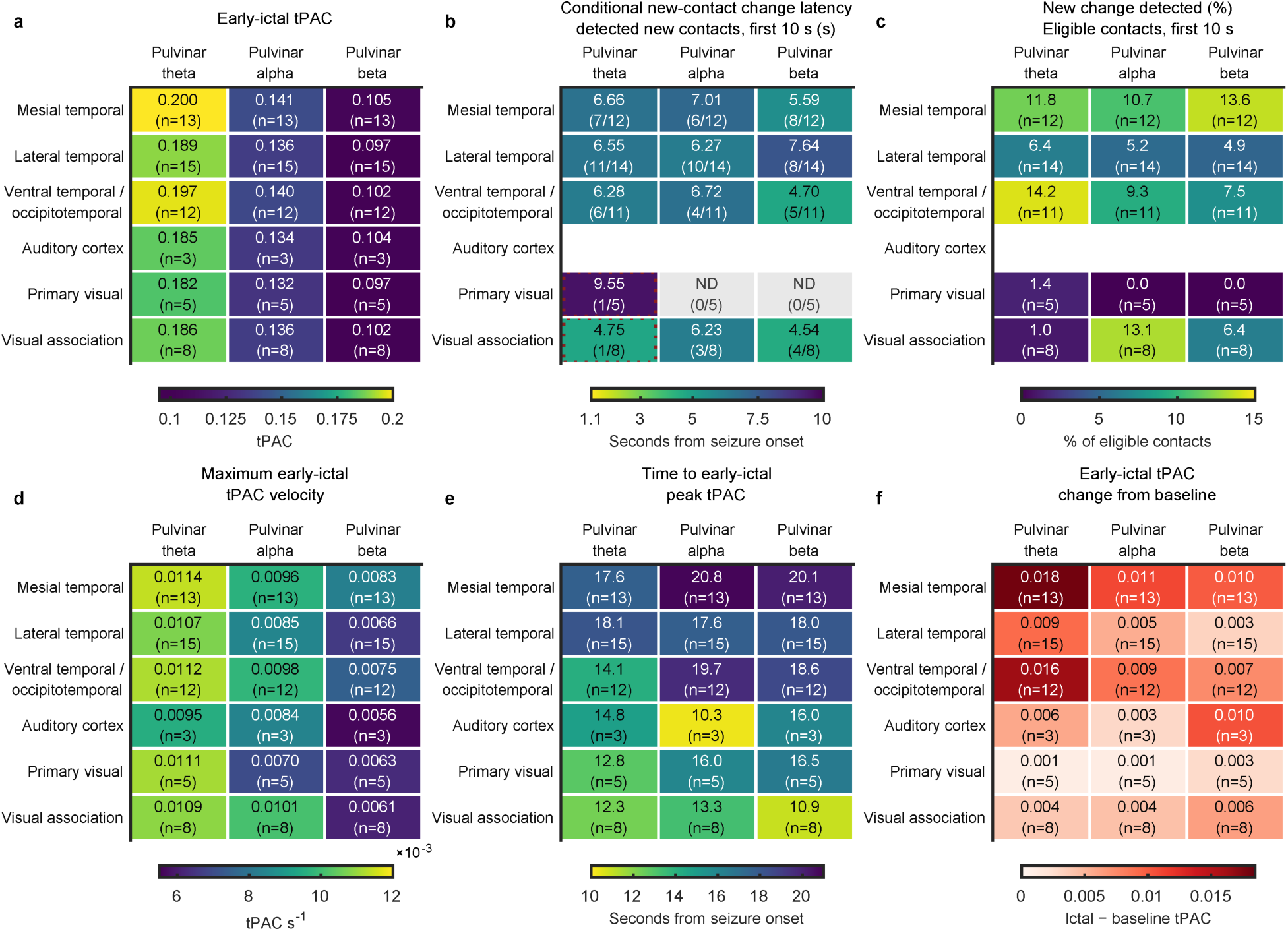
Early-ictal pulvinar-cortical coupling in temporal and occipital cortical subgroups. Heat maps showing regional coupling magnitude and temporal characteristics across predefined temporal and occipital cortical subgroups for pulvinar theta-, alpha-, and beta-phase coupling. **a,** Mean tPAC during the early ictal third. **b,** Conditional contact-level detection latency among contacts showing a new coupling change within 10 s after electrographic seizure onset. **c,** Proportion of eligible contacts with a new peri-onset detection within 10 s. **d,** Maximum tPAC velocity during the early ictal third, calculated from the nominally 10-s smoothed trajectory (tPAC s⁻¹). **e,** Time from seizure onset to peak tPAC within the early ictal third (s). **f,** Absolute early-ictal-minus-baseline tPAC difference (dimensionless). Temporal subgroups included mesial temporal, lateral temporal neocortical, ventral temporal/occipitotemporal, and auditory cortex; occipital subgroups included primary visual (calcarine/cuneus) and visual association regions. For **b** and **c**, detections were classified as new only in the absence of a recent pre-onset detection (−5 to 0 s). Cell labels indicate values and contributing participant numbers (responding/eligible participants in **b**); white cells indicate fewer than three eligible participants, ND indicates no detection, and dotted outlines indicate fewer than three responding participants. These measures are descriptive and do not establish a sequence of cortical recruitment.

Focused peri-onset analysis identified the earliest regional departure from the preictal reference in pulvinar beta-phase coupling with temporal cortex (7.70 s). Contact-level analyses further revealed regional variability in peri-onset coupling changes (Fig. 7b, c), with the highest detection fractions in ventral temporal/occipitotemporal cortex for theta-phase coupling, visual association cortex for alpha-phase coupling, and mesial temporal cortex for beta-phase coupling. Conditional detection latencies varied across regions without a consistent anatomical sequence.

Regional temporal analyses revealed further heterogeneity in pulvinar-cortical coupling dynamics. Occipital cortex showed the shortest time to peak tPAC within the early ictal third across all three pulvinar phase bands (12.8, 13.1, and 12.1 s for theta, alpha, and beta, respectively; n = 8; subgroup values in Fig. 7e). Maximum tPAC velocity also varied by region, with the highest values in temporal cortex for theta (0.0115 tPAC s⁻¹), occipital cortex for alpha (0.0102 tPAC s⁻¹), and parietal cortex for beta (0.0063 tPAC s⁻¹; subgroup values in Fig. 7d). Regional differences in maximum tPAC velocity were descriptive and sensitive to smoothing parameters.

### 3.6 Pulvinar subdivision-specific coupling patterns

Pulvinar subdivision coverage was unequal, with PuM represented in 13 participants, PuL in four, and PuA in two (Fig. 8a). Across cortical targets, mean early-ictal tPAC was similar between PuM and PuL for theta-(0.1875 versus 0.1871), alpha-(0.1346 versus 0.1345), and beta-phase coupling (0.0983 versus 0.0993; Fig. 8b). Whole-ictal baseline-to-peak secant rates were 0.86, 0.53, and 0.44 × 10⁻³ tPAC s⁻¹ for PuM and 1.00, 0.69, and 0.56 × 10⁻³ tPAC s⁻¹ for PuL across theta-, alpha-, and beta-phase coupling, respectively (Fig. 8c). PuA estimates were exploratory because of limited sampling, and subdivision comparisons were descriptive.

**Figure 8.**
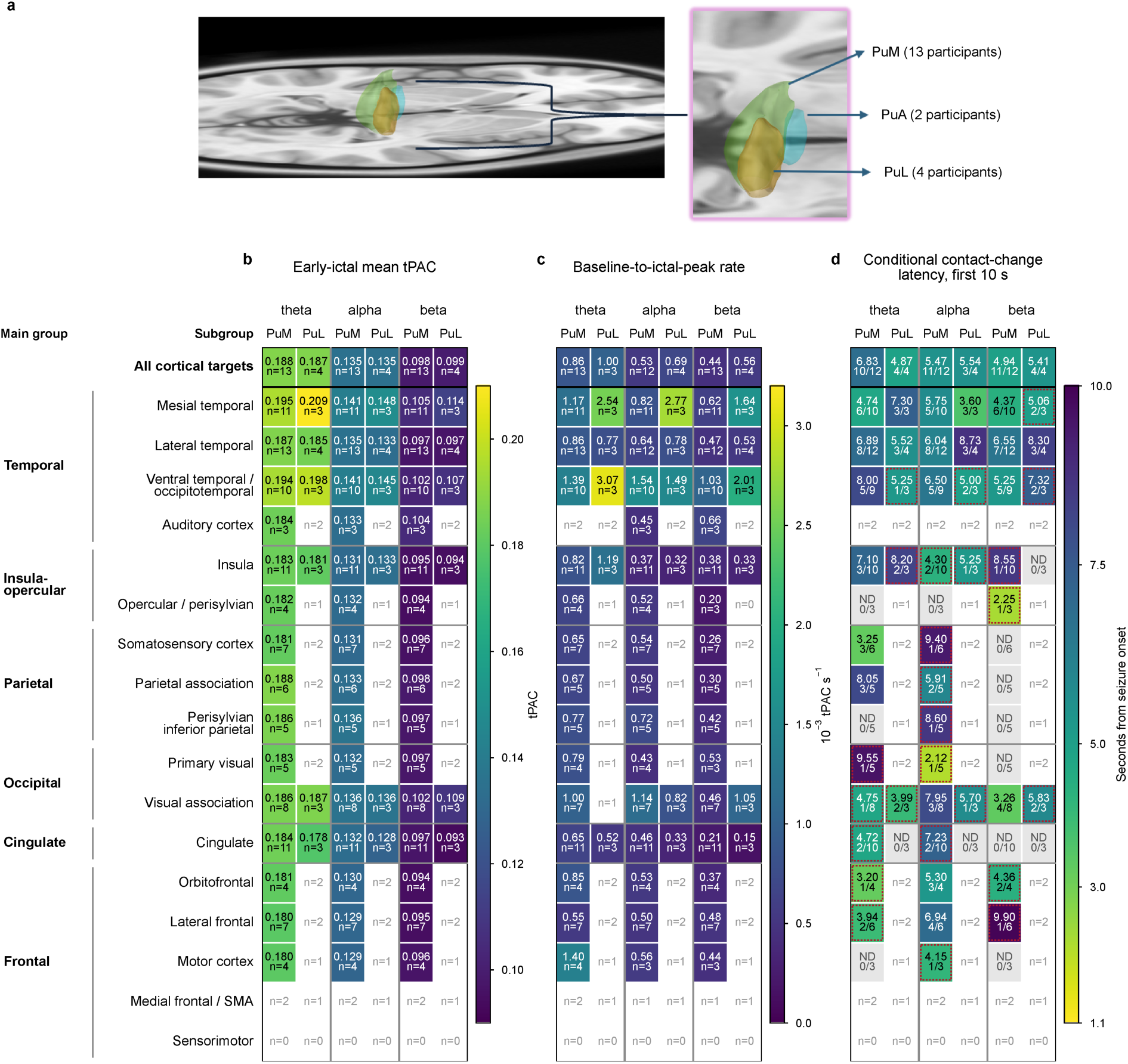
Pulvinar subdivision-specific patterns of pulvinar-cortical coupling. **a**, Distribution of sampled pulvinar subdivisions across participants (PuM, n = 13; PuL, n = 4; PuA, n = 2). **b,** Mean early-ictal pulvinar theta-, alpha-, and beta-phase tPAC across cortical targets for PuM and PuL. **c,** Whole-ictal baseline-to-peak tPAC secant rates across the three phase bands for PuM and PuL. **d,** Conditional peri-onset coupling detection latencies across cortical regions and phase bands, stratified by pulvinar subdivision. Subdivision comparisons were descriptive because of unequal sampling, and PuA was excluded from **b-d** because only two participants contributed data.

Within PuM, early-ictal coupling was numerically greatest in parahippocampal cortex for theta- and alpha-phase coupling (0.1965 and 0.1439 tPAC; n = 9) and in inferior occipital cortex for beta-phase coupling (0.1081 tPAC; n = 4). Within PuL, temporal targets showed the greatest theta- and alpha-phase coupling, whereas occipital targets showed the greatest beta-phase coupling.

Subdivision-specific peri-onset detection latencies varied across cortical regions and phase bands without a consistent regional sequence (Fig. 8d). Overall, the available data did not establish a reproducible advantage of any pulvinar subdivision or determine whether PuM predominance contributed to the broader regional coupling patterns. Because eligible pulvinar seeds were averaged within cortical targets before the focused peri-onset analysis, subdivision-specific estimates of earliest baseline departure were unavailable.

## 4. Discussion

Direct pulvinar recordings obtained during human stereo-EEG monitoring allowed us to characterize pulvinar involvement in seizure networks through ictal recruitment patterns and time-resolved pulvinar-cortical coupling. Four principal findings emerged. First, electrophysiological assessment demonstrated that pulvinar recruitment frequently occurred at or near cortical seizure onset, indicating early thalamic involvement in sampled seizure networks.

Second, pulvinar-cortical coupling was preferentially greater in SOZ cortex across pulvinar theta-, alpha-, and beta-phase frequencies, with greater seizure-related increases in SOZ cortex for alpha- and beta-phase coupling. Third, coupling exhibited distinct temporal and frequency- dependent dynamics, with pulvinar beta-phase coupling in SOZ cortex showing the earliest cohort-level departure from the preictal baseline, greater post-onset rise rates across all three phase bands, and earlier ictal maxima for theta- and alpha-phase coupling compared with non-SOZ cortex. Fourth, coupling exhibited anatomical heterogeneity, with numerically greater early-ictal theta- and alpha-phase coupling in temporal regions, particularly parahippocampal cortex, and beta-phase coupling in occipital regions, particularly inferior occipital cortex. Together, these findings demonstrate that the pulvinar is dynamically engaged during human seizures, with coupling patterns that vary across cortical networks, frequency bands, and seizure evolution, providing a physiological framework for investigating its role in seizure networks and its potential relevance as a target for neuromodulation in epilepsy.

Pulvinar recruitment in our cohort coincided with electrographic cortical onset in more than half of seizures and occurred within 10 s in approximately three quarters of events. Immediate pulvinar recruitment was more frequent among seizures with a cortical low-voltage fast activity (LVFA) onset pattern, suggesting an association between cortical onset morphology and the timing of pulvinar involvement. Together, these observations indicate that the pulvinar can become engaged very early during seizure evolution, including near-simultaneously with electrographic cortical onset, highlighting its early participation in the sampled seizure networks. These observations are consistent with prior stereo-EEG studies reporting ictal activity in the medial pulvinar in most temporal lobe seizures^16^, thalamic involvement in most patients with focal epilepsy^18^, and earlier or more prominent involvement of the pulvinar than of the anterior nucleus in many patients^13,14^.

Time-resolved phase-amplitude coupling (tPAC) provides a framework for characterizing the temporal evolution of thalamocortical interactions during seizures. Conventional PAC estimators, including mean-vector-length and entropy-based modulation-index approaches, quantify coupling over defined analysis windows^21,22^. Shortening these windows improves temporal resolution but can compromise estimation reliability, motivating approaches designed to track coupling dynamics at shorter timescales^28^. Using tPAC, we identified preferential pulvinar-cortical coupling with clinically defined seizure-onset-zone (SOZ) cortex. Absolute ictal tPAC was greater in SOZ than non-SOZ contacts across all three pulvinar phase-frequency bands, while seizure-related increases from baseline were greater in SOZ cortex for alpha- and beta-phase coupling. Exploratory classification further suggested that theta-phase tPAC features may help distinguish SOZ from non-SOZ contacts in held-out participants, although differences between pooled and within-participant performance warrant further evaluation. These findings suggest that pulvinar-cortical interactions are not uniformly distributed across the sampled cortical network but are more strongly expressed within SOZ regions.

Frequency-specific coupling has been described in both cortical and thalamic recordings. Coupling involving delta-, theta-, and alpha-phase and gamma/ripple amplitudes has been observed in temporal lobe epilepsy^30^, and theta-phase modulation of ripple activity has been reported during focal seizures^53^. Separately, theta-phase (4-6 Hz) coupling to 80-150-Hz activity has been recorded at rest in the human anterior thalamic nuclei^54^, while pulvinar interactions with visual cortex during visuospatial attention in macaques were prominent in the alpha range (8-15 Hz^55^). Against this physiological background, our analyses revealed distinct temporal characteristics of pulvinar-cortical coupling across phase-frequency bands. Post-onset rise rates were greater in SOZ than non-SOZ cortex across all three bands, while ictal coupling reached its maximum approximately 10 s earlier for theta-phase and 8 s earlier for alpha-phase coupling in SOZ cortex. Exploratory duration-normalized analyses further suggested time-local SOZ-non-SOZ differences, with the greatest theta-phase separation near the middle of seizures and greater alpha- and beta-phase baseline-relative coupling in SOZ cortex during the early ictal period. Although these exploratory temporal findings were sensitive to multiple-comparison correction and localization criteria, they suggest that SOZ-associated pulvinar-cortical coupling may differ in both magnitude and temporal evolution across seizure stages.

Focused peri-onset analyses provided additional insights into pulvinar-cortical coupling dynamics around electrographic seizure onset, a period relevant to early seizure detection and therapeutic intervention. Cohort-mean pulvinar-cortical coupling in SOZ cortex was elevated during the recent preictal period relative to the distant preictal reference across all three phase bands, suggesting that coupling changes may precede electrographic seizure onset, although these effects did not survive participant-level correction (Supplementary Results). Following seizure onset, pulvinar beta-phase coupling in SOZ cortex showed the earliest cohort-level departure from the preictal reference at the first evaluable endpoint (1.65 s), followed by alpha-phase coupling at 3.85 s, while theta-phase coupling and all three bands in non-SOZ cortex showed no corresponding departure within the evaluated interval. The baseline-referenced SOZ-non-SOZ difference in beta-phase coupling also departed at 1.65 s, followed by alpha-phase coupling at 9.35 s. Complementary contact-level analyses showed a greater proportion of SOZ than non-SOZ contacts with newly detected beta-phase coupling changes within 10 s of seizure onset (21.3% versus 4.5%), although this difference was sensitive to cohort selection (Supplementary Results). The early beta-phase departure persisted after accounting for recent preictal evolution, supporting an additional post-onset change beyond the preceding coupling trajectory. Together, these findings suggest that early pulvinar beta-phase coupling may reflect SOZ-associated network engagement, while participant heterogeneity and the absence of significant direct post-onset SOZ-non-SOZ contrasts after correction limit conclusions about its consistency across individuals or precise biological onset.

Meta-analysis of functional neuroimaging identifies widespread but regionally differentiated pulvinar coactivation^6^, human diffusion tractography supports a putative pulvinar-amygdala pathway^56^, and single-pulse stimulation in epilepsy patients demonstrates evoked connectivity between the pulvinar and mesial temporal and distributed cortical structures^15,57,58^. Consistent with this anatomical organization, our findings revealed regional variation in the spectral and temporal characteristics of pulvinar-cortical coupling during seizures. During the early ictal period, temporal cortex exhibited the numerically greatest theta- and alpha-phase coupling (0.1924 and 0.1372 tPAC), particularly in parahippocampal regions, whereas occipital cortex numerically showed the greatest beta-phase coupling (0.1012 tPAC), especially within inferior regions. Temporal alpha-phase coupling was significantly greater than parietal and frontal coupling in matched comparisons (P = 0.0044 and 0.031, respectively). Mesial temporal coupling was also numerically greater than lateral temporal neocortical coupling across all three bands. These findings suggest that frequency-specific pulvinar-cortical interactions may reflect the anatomical organization of distributed seizure networks.

Regional differences extended to baseline-relative changes and temporal dynamics. Ventral temporal/occipitotemporal cortex showed a relatively large early-ictal increase in theta-phase coupling (absolute ictal-minus-baseline tPAC difference of 0.016) and the highest proportion of newly detected peri-onset theta-phase coupling changes. Pulvinar beta-phase coupling with temporal cortex was the only absolute broad-region trajectory to depart from its corrected preictal reference within the first 9.9 s (at 7.70 s), whereas occipital cortex showed the shortest times to peak coupling within the early ictal third across theta, alpha, and beta bands (12.8, 13.1, and 12.1 s). These observations suggest that coupling magnitude, early detectability, and peak timing capture distinct aspects of regional pulvinar involvement during seizures.

Evoked-potential mapping across pulvinar subregions has demonstrated preferential cortical responses to stimulation of the lateral, ventromedial, and dorsomedial pulvinar in visual, lateral temporal, and parietal cortices, respectively^59^. Subdivision sampling in our cohort was unequal (PuM, n = 13; PuL, n = 4; PuA, n = 2). Within PuM, parahippocampal cortex showed prominent theta- and alpha-phase coupling (0.1965 and 0.1439 tPAC), consistent with the broader regional findings. Limited PuL and PuA sampling restricted subdivision-specific comparisons and the generalizability of these temporal-network patterns. Together, these findings suggest that pulvinar-cortical coupling exhibits regionally differentiated spectral and temporal dynamics, potentially reflecting the organization of thalamocortical networks rather than a spatially uniform ictal response.

Several limitations should be considered. This retrospective study included 15 participants undergoing clinically indicated sEEG, resulting in heterogeneous cortical and pulvinar sampling, particularly across pulvinar subnuclei. Variation in seizure-onset networks further constrained network-specific inference, while clinically defined SOZ boundaries were dependent on available electrode coverage. Primary statistical inference was performed at the participant level to account for repeated seizures, whereas seizure-level analyses of visual recruitment and clinical timing were considered descriptive. The temporal resolution of tPAC estimation (1.1-s windows) and offline processing limit interpretation of peri-onset departures as precise biological coupling onset times or real-time latencies; similarly, time to peak reflects maximal coupling rather than the onset of pulvinar recruitment. Some frequency- and time-specific findings were sensitive to anatomical localization criteria or multiple-comparison correction and should therefore be considered exploratory. The SOZ classification analysis was evaluated using leave-one-participant-out validation within the present cohort and requires independent validation to establish generalizability. Finally, tPAC reflects statistical dependence rather than directional or causal interactions, and may be influenced by ictal changes in spectral power and waveform morphology rather than exclusively reflecting mechanistic cross-frequency coupling^20,26^.

Pulvinar neuromodulation has shown promising preliminary clinical outcomes in small studies of posterior cortical, temporal, and other focal epilepsies, although evidence remains limited and optimal approaches to patient selection and stimulation targeting are not established^7,10,11^. Our findings suggest that the relevance of the pulvinar as a neuromodulation target may depend on how strongly and dynamically it is embedded within an individual patient’s seizure network. Although our study did not evaluate neuromodulation outcomes, the observed variability in pulvinar recruitment, preferential coupling with SOZ cortex, and frequency-dependent temporal dynamics have potential implications for pulvinar-targeted neuromodulation. These findings raise important questions regarding patient and seizure-network selection, optimal stimulation targets, and the timing of stimulation relative to seizure evolution. Dynamic pulvinar-cortical coupling may offer complementary information about ictal network state beyond that captured by conventional electrographic detection features, such as line length, band-limited power, and area-based measures^60,61^. Addressing these questions will require prospective studies evaluating whether pulvinar-cortical network characteristics can inform patient selection, stimulation targeting, and real-time detection strategies, and how these features relate to stimulation parameters and clinical outcomes.

## Data availability statement

Data will be made available upon reasonable request and with data agreements.

## Author statement

We confirm that we have read the Journal’s position on issues involved in ethical publication and affirm that this report is consistent with those guidelines.

## Author contributions

Y.C.: Data curation, Formal analysis, Methodology, Software, Visualization, Writing - original draft, Writing - review & editing. C.G.: Methodology, Visualization, Writing - review & editing. S.H., G.G., and D.S.: Investigation (stereo-EEG implantation), Writing - review & editing. S.A.: Conceptualization, Data curation, Formal analysis, Methodology, Software, Supervision, Writing - original draft, Writing - review & editing.

## Ethics statement

IRB-waived informed consent

## Supporting information

Supplementary Results

## SUPPLEMENTARY

### Supplementary Results

#### Recent-preictal and contact-level sensitivity analyses of peri-onset pulvinar-cortical coupling

Cohort-mean SOZ tPAC was elevated during recent-preictal intervals relative to the distant reference by 0.255 reference SD for beta (baseline-family-adjusted P = 0.0002), 0.209 SD for alpha (P = 0.0004), and 0.166 SD for theta (P = 0.0432). These effects did not survive participant-family correction, and no significant recent-preictal slope or reference-drift effect was identified after correction. Following subtraction of linear forecasts fitted to recent pre-onset activity, both the SOZ beta residual and direct SOZ-non-SOZ beta residual departed from their corrected baseline envelopes at 1.65 s (baseline-family-adjusted P = 0.0246 for each), whereas alpha and theta did not. Both beta departures persisted at the first evaluable endpoint (3.85 s) under the more conservative phase-support restriction. However, no local-linear participant-level test survived family-wise correction, indicating a cohort-level beta departure without corrected evidence of a consistent participant-level response.

The participant-level conclusion was sensitive to the multiplicity-correction family. Under the prespecified distant-reference analysis, correction across nine trajectories (six SOZ/non-SOZ phase-band trajectories and three direct SOZ-non-SOZ contrasts) and 34 pre- and post-onset endpoints yielded a minimum post-onset adjusted P = 0.251 for the direct beta contrast at 3.85 s. A post hoc analysis restricted to the three direct SOZ-non-SOZ contrasts across 16 post-onset endpoints yielded an adjusted P = 0.033 at the same endpoint. The broader prespecified family was retained for primary inference.

In the complementary contact-level analysis, the participant-balanced beta detection fraction by 10 s was greater in SOZ than non-SOZ cortex (21.3% versus 4.5%; mean participant-paired difference, 16.8 percentage points; 95% participant-bootstrap interval, 6.5-30.3; Holm-adjusted P = 0.021; 14 participants, 28 seizures). The 2-and 5-s analyses included 14 participants and 29 seizures and were nonsignificant after correction. One additional seizure was excluded from the 10-s analysis because it lacked complete evaluable follow-up through that endpoint.

The 10-s finding was sensitive to cohort selection, with adjusted P = 0.064 when including all available baselines (15 participants, 29 seizures) and P = 0.504 under stricter pulvinar seed-localization criteria (11 participants, 21 seizures). These findings suggest an excess of early beta detections in SOZ cortex that was sensitive to cohort selection. Estimator temporal resolution, participant heterogeneity, and differences in inferential endpoints preclude conclusions regarding precise biological coupling onset or the sequence of cortical recruitment.

**Supplementary Figure 1.**
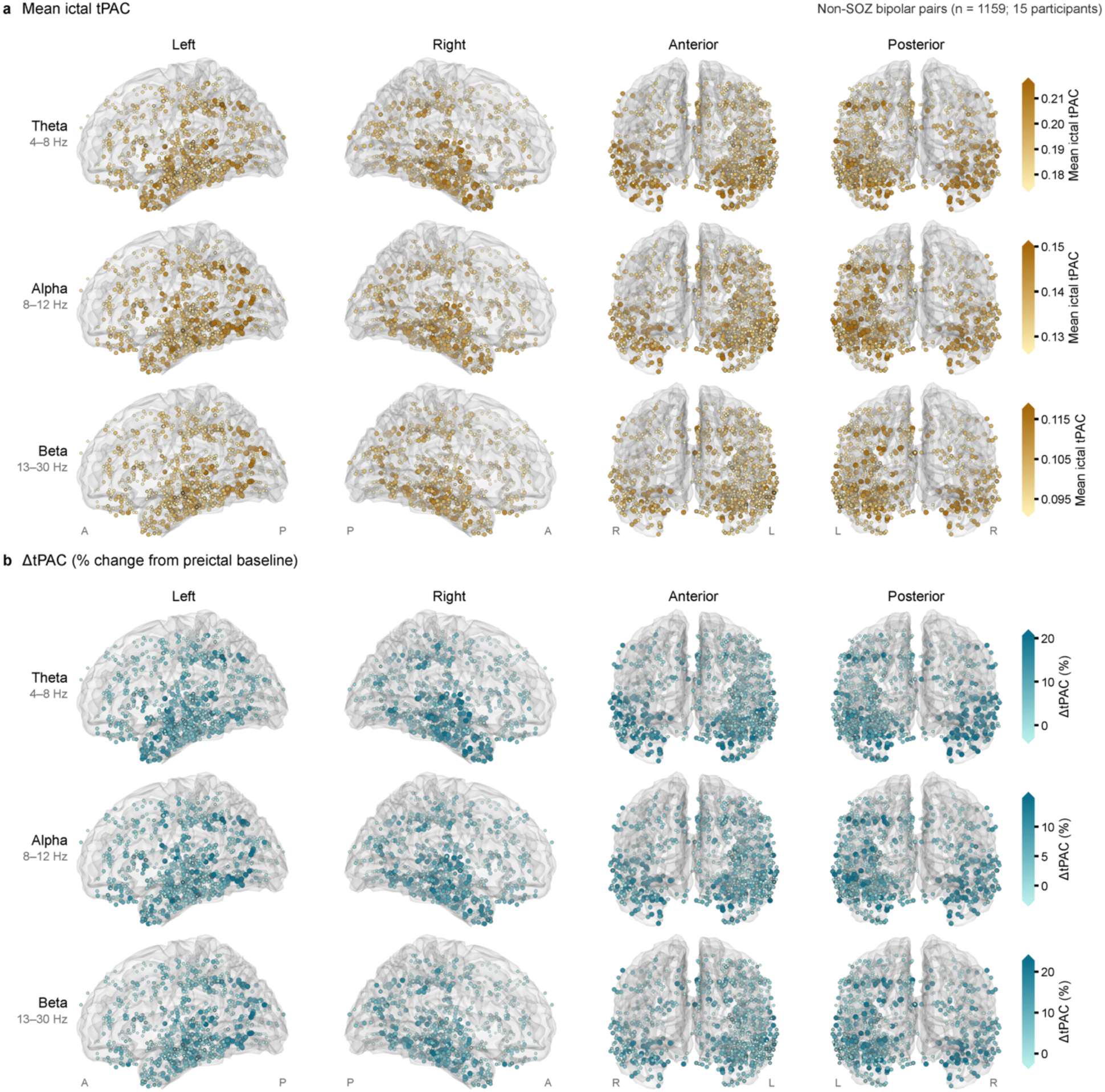
Spatial distribution of mean ictal tPAC and ΔtPAC across sampled non-SOZ cortex. Glass-brain maps of 1,159 non-seizure-onset-zone (non-SOZ) cortical bipolar targets from 15 participants, shown as in Fig. 6 in left, right, anterior, and posterior views for pulvinar theta-, alpha-, and beta-phase coupling. **a,** Mean ictal tPAC over the full electrographic seizure interval. **b,** Baseline-relative change in tPAC (ΔtPAC), expressed as percentage change from the preictal baseline (−100 to −10 s). Marker placement, projection, and scaling are as in Fig. 6. Within each band and measure, color limits (5th-95th percentiles of all 1,308 mapped targets) are identical to those used for the SOZ maps, allowing direct visual comparison between figures. A, anterior; P, posterior; L, left; R, right. These maps are descriptive and were not used for statistical inference.

**Supplementary Table 1.**
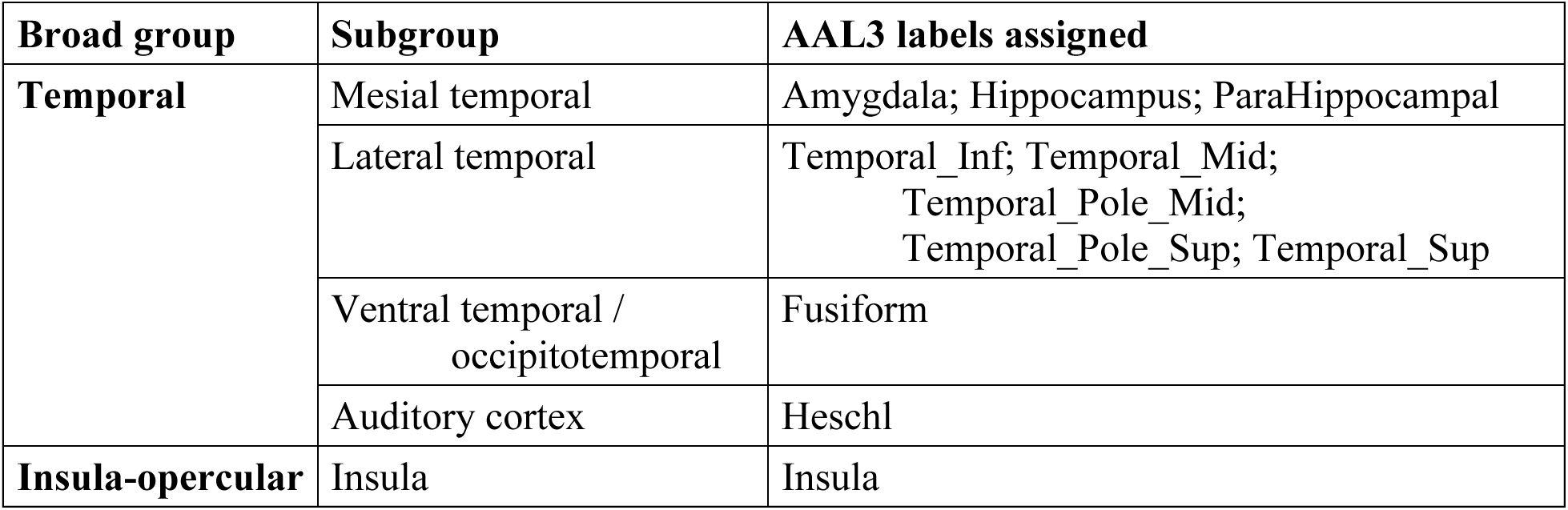
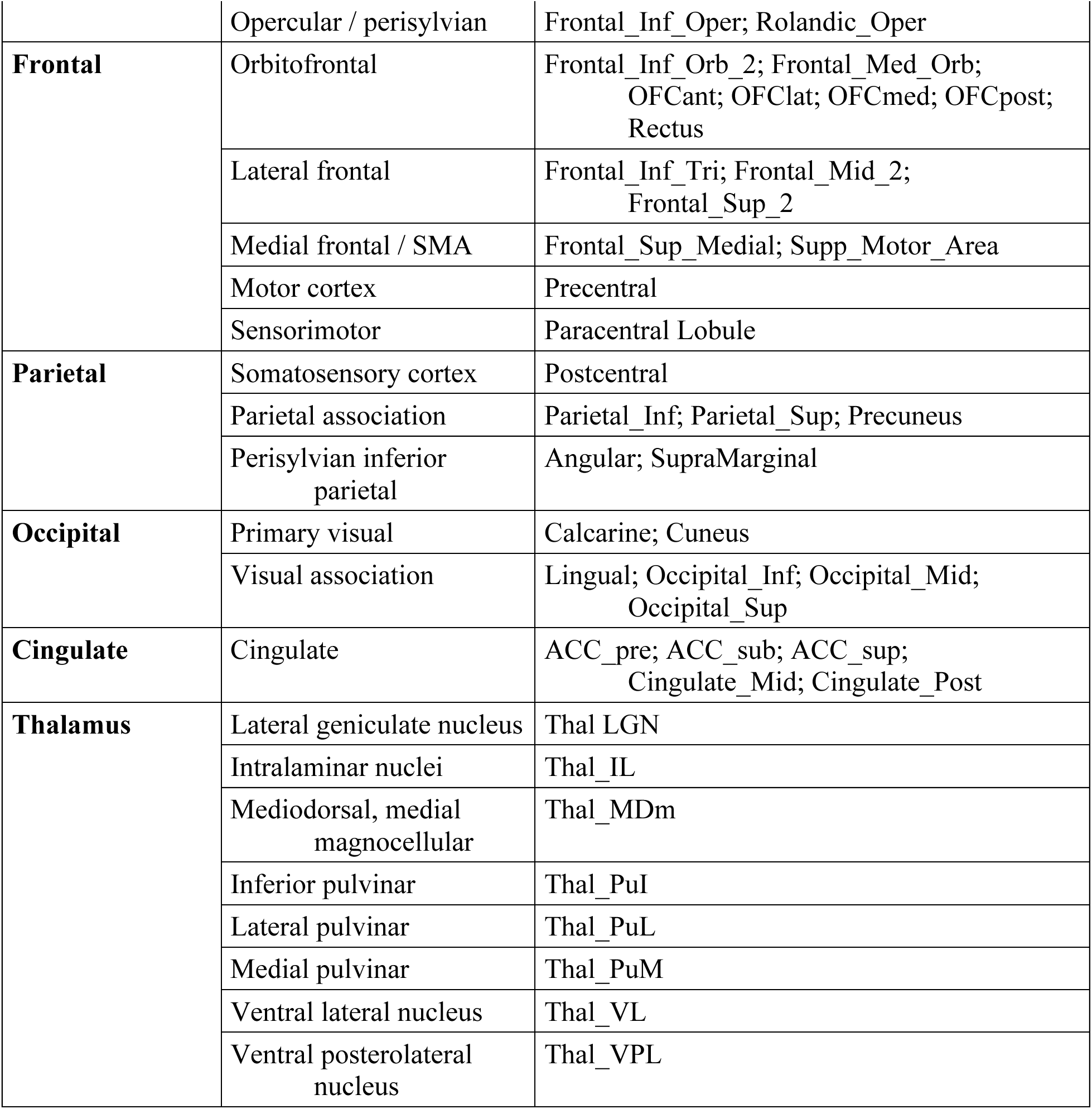
Cortical anatomical grouping scheme used for regional analyses. Broad cortical groups and corresponding finer anatomical subgroups used to classify bipolar cortical targets for regional pulvinar-cortical coupling analyses.

