## Supplementary Results for "Pulvinar-cortical coupling in human epilepsy: spatiotemporal seizure dynamics with implications for neuromodulation"

| **Broad group** | **Subgroup** | **AAL3 labels assigned** |
|---|---|---|
| **Temporal** | Mesial temporal | Amygdala; Hippocampus; ParaHippocampal |
| | Lateral temporal | Temporal_Inf; Temporal_Mid; Temporal_Pole_Mid; Temporal_Pole_Sup; Temporal_Sup |
| | Ventral temporal / occipitotemporal | Fusiform |
| | Auditory cortex | Heschl |
| **Insula-opercular** | Insula | Insula |

| Broad group | Subgroup | AAL3 labels assigned |
|---|---|---|
| | Opercular / perisylvian | Frontal_Inf_Oper; Rolandic_Oper |
| **Frontal** | Orbitofrontal | Frontal_Inf_Orb_2; Frontal_Med_Orb; OFCant; OFClat; OFCmed; OFCpost; Rectus |
| | Lateral frontal | Frontal_Inf_Tri; Frontal_Mid_2; Frontal_Sup_2 |
| | Medial frontal / SMA | Frontal_Sup_Medial; Supp_Motor_Area |
| | Motor cortex | Precentral |
| | Sensorimotor | Paracentral Lobule |
| **Parietal** | Somatosensory cortex | Postcentral |
| | Parietal association | Parietal_Inf; Parietal_Sup; Precuneus |
| | Perisylvian inferior parietal | Angular; SupraMarginal |
| **Occipital** | Primary visual | Calcarine; Cuneus |
| | Visual association | Lingual; Occipital_Inf; Occipital_Mid; Occipital_Sup |
| **Cingulate** | Cingulate | ACC_pre; ACC_sub; ACC_sup; Cingulate_Mid; Cingulate_Post |
| **Thalamus** | Lateral geniculate nucleus | Thal LGN |
| | Intralaminar nuclei | Thal_IL |
| | Mediodorsal, medial magnocellular | Thal_MDm |
| | Inferior pulvinar | Thal_PuI |
| | Lateral pulvinar | Thal_PuL |
| | Medial pulvinar | Thal_PuM |
| | Ventral lateral nucleus | Thal_VL |
| | Ventral posterolateral nucleus | Thal_VPL |

**Supplementary Table 1 |** Cortical anatomical grouping scheme used for regional analyses. Broad cortical groups and corresponding finer anatomical subgroups used to classify bipolar cortical targets for regional pulvinar-cortical coupling analyses.

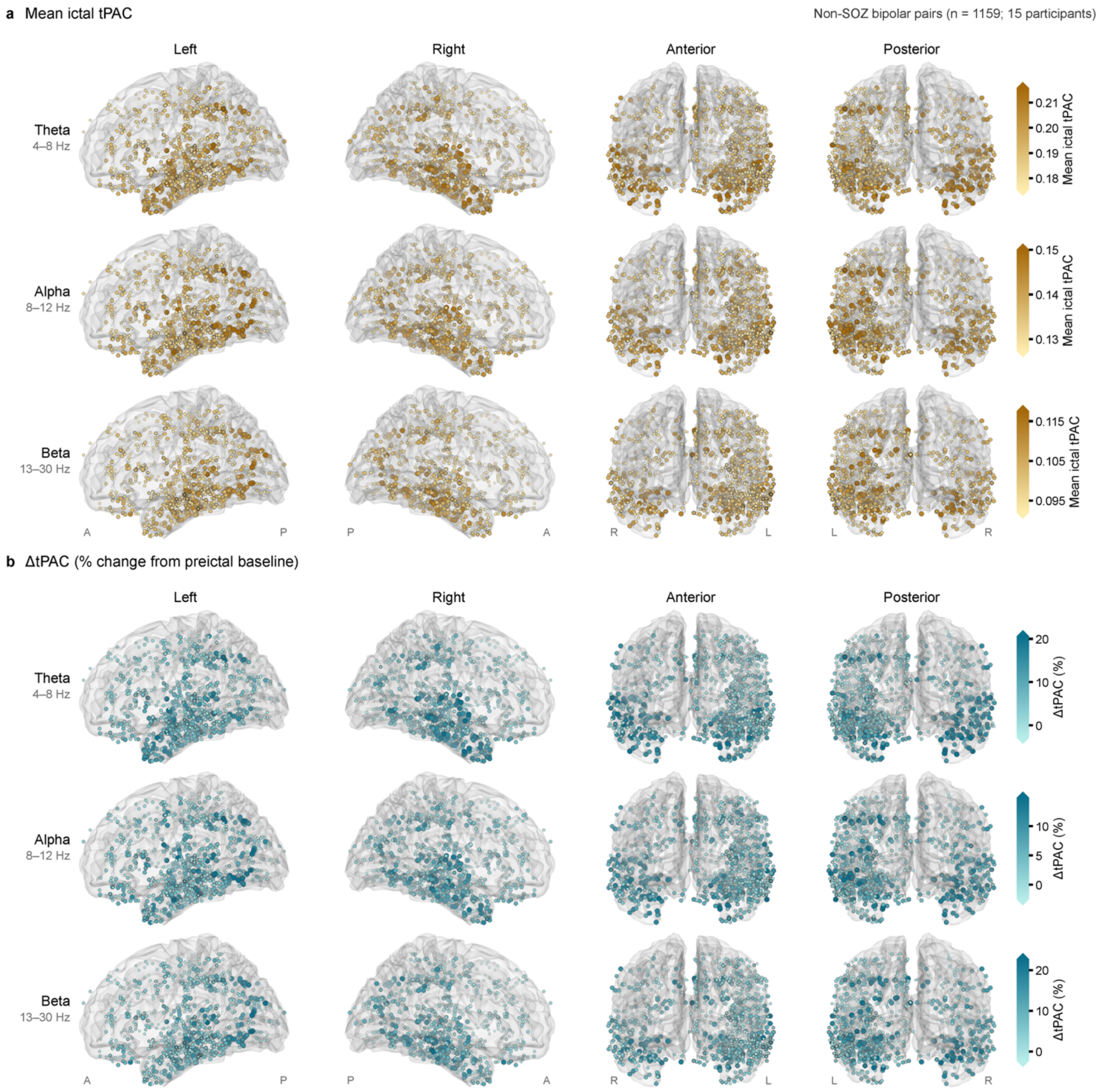


**Supplementary Figure 1 | Spatial distribution of mean ictal tPAC and ΔtPAC across sampled non-SOZ cortex**

Glass-brain maps of 1,159 non-seizure-onset-zone (non-SOZ) cortical bipolar targets from 15 participants, shown as in Fig. 6 in left, right, anterior, and posterior views for pulvinar theta-, alpha-, and beta-phase coupling. **a,** Mean ictal tPAC over the full electrographic seizure interval. **b,** Baseline-relative change in tPAC (ΔtPAC), expressed as percentage change from the preictal baseline (−100 to −10 s). Marker placement, projection, and scaling are as in Fig. 6. Within each band and measure, color limits (5th-95th percentiles of all 1,308 mapped targets) are identical to those used for the SOZ maps, allowing direct visual comparison between figures. A, anterior; P, posterior; L, left; R, right. These maps are descriptive and were not used for statistical inference.
